# Ecdysone coordinates plastic growth with robust pattern in the developing wing

**DOI:** 10.1101/2020.12.16.423141

**Authors:** André Nogueira Alves, Marisa Mateus Oliveira, Takashi Koyama, Alexander Shingleton, Christen Mirth

## Abstract

Animals develop in unpredictable, variable environments. In response to environmental change some aspects of development adjust to generate plastic phenotypes. Other aspects of development, however, are buffered against environmental change to produce robust phenotypes. How organ development is coordinated to accommodate both plastic and robust developmental responses is poorly understood. Here, we demonstrate that the steroid hormone ecdysone coordinates both plasticity of organ size and robustness of organ pattern in the developing wings of the fruit fly *Drosophila melanogaster*. Using fed and starved larvae that lack prothoracic glands, which synthesise ecdysone, we show that nutrition regulates growth both via ecdysone and via an ecdysone-independent mechanism, while nutrition regulates patterning only via ecdysone. We then demonstrate that growth shows a graded response to ecdysone concentration, while patterning shows a threshold response. Collectively, these data support a model where nutritionally-regulated ecdysone fluctuations confer plasticity by regulating disc growth in response to basal ecdysone levels, and confers robustness by initiating patterning only once ecdysone peaks exceeds a threshold concentration. This could represent a generalizable mechanism through which hormones coordinate plastic growth with robust patterning in the face of environmental change.

## Introduction

Developing animals respond to changes in their environment in a multitude of ways, for example, altering how long and how fast they grow, the time it takes them to mature, and their reproductive output [1, 2]. Other aspects of their phenotype, however, must be unresponsive to environmental change to ensure that they function correctly regardless of environmental conditions. This presents a particular problem for morphological traits of developing animals. For any given trait, some aspects, such as final organ size, vary with changes in the environment, a phenomenon termed plasticity [3–7]. Other aspects, like patterning the cell types within an organ necessary for it to function, remain constant across environmental conditions, and are thus termed robust [6–9]. For many organs, growth and patterning occur at the same time during development, and may even be regulated by the same hormones [6]. How then do organs achieve plasticity in size while maintaining robustness of pattern?

If we want to extract general principles of how organisms regulate their development in variable environments, we need to understand how developmental processes unfold over time. Several recent studies that have applied systems approaches to development offer excellent examples, frequently employing methods to quantify how gene expression patterns change over time. These studies have used the dynamic changes in expression patterns to uncover the rules governing how insects build their segments [10, 11], how the gene regulatory network underlying segmentation evolves [12–17], how morphogen gradients scale across organs and bodies [18–23], how sensory organs are positioned within epithelia [24], and how somites and digits form in vertebrates [25–27]. The power of these approaches is that they provide a framework for understanding how genes interact within a network to generate pattern that can be applied across a variety of contexts.

The success of these studies is, in part, due to the fact that the gene regulatory networks underlying each of these processes have been well described in their respective developmental contexts. In contrast, the gene regulatory networks governing growth and patterning at later stages of development, even at later stages of embryonic development, are not as well resolved. If we further complicate this by comparing development across environmental conditions and even across traits, approaches that rely on understanding the configuration of gene regulatory networks become much more difficult to implement.

Nevertheless, we can still use the principle of comparing the dynamics of developmental processes across environments to gain useful insights into the relationship between plasticity and robustness. Many types of environmental conditions impact organ development to induce changes in body and organ size. Malnutrition or starvation reduces growth rates in all animals, resulting in smaller body and organ sizes [28–30]. Similarly, changing temperature can alter animal growth. In insect species, rearing animals in warmer conditions results in smaller adult body sizes when compared to animals reared under cooler conditions [31–38]. Other factors like oxygen availability and the presence of toxic or noxious compounds also act to alter animal sizes [39–41]. Examining how organ growth and patterning progress across these environmental conditions helps us to understand how these two processes are coordinated.

We already have some understanding of the mechanisms that regulate growth and patterning in response to changing environmental conditions. The genetic mechanisms underlying plasticity in growth is best elucidated in insects. In insects, changes in available nutrition affect the synthesis and secretion of the conserved insulin-like peptides [43–45]. Insulin-like peptides bind to the Insulin Receptor in target tissues and activate the insulin signalling cascade, ultimately leading to increased growth [44, 46, 47]. Starvation reduces the concentration of insulin-like peptides in the bloodstream, and the resulting decrease in insulin signalling causes organs to grow more slowly [45, 48].

While changes in insulin signalling are known to affect organ size, they have little effect on organ pattern [49]. However, studies in the fruit fly *Drosophila melanogaster* have shown that, at least in this insect, insulin acts to control the synthesis of a second developmental hormone, the steroid hormone ecdysone [50–53]. Most of the body and organ growth in *D. melanogaster* occurs in the third, and final, larval instar, after which the animal initiates metamorphosis at pupariation. Either starving or reducing insulin signalling early in the third instar delays the timing of ecdysone synthesis, thereby prolonging the length of the third instar and the time it takes to metamorphose [50–54].

In addition to its effects on developmental time, ecdysone controls the growth of the developing adult organs [55–58]. In *D. melanogaster* larvae, many of the adult organs form and grow inside the larvae as pouches of cells called imaginal discs. If ecdysone synthesis is reduced or if the glands that produce ecdysone, the prothoracic glands (PG), are ablated, these imaginal discs grow at greatly reduced rates [56, 59].

Ecdysone signalling also regulates organ patterning. Reducing ecdysone signalling in either the wing imaginal disc or the developing ovary causes substantial delays in their patterning [56, 59–61]. In the wing disc, reducing ecdysone signalling stalls the progression of patterning of sensory bristles [56, 59]. Similarly, in the ovary terminal filament cell specification and the rate of terminal filament addition both require ecdysone to progress normally [60, 61]. Given its role in both the patterning and the growth of imaginal discs and ovaries, ecdysone is potentially a key coordinator of plastic growth and robust pattern.

Characterising organ growth rates is experimentally straight forward, requiring only accurate measurement of changes in organ size over time. To quantify the progression of organ patterning, however, requires developing a staging scheme. We previously developed such a scheme for the wing imaginal disc in *D. melanogaster*. This scheme makes use of the dynamic changes in expression from the moult to third instar to pupariation of up to seven patterning gene products in the developing wing [42]. Two of these patterning gene products, Achaete and Senseless, can be classed into seven different stages throughout third instar development ([42], Figure 1A), providing us with the ability to quantify the progression of wing disc pattern over a variety of conditions. In short, by describing patterning on a near continuous scale, our scheme not only allows us to determine under what conditions patterning is initiated, but also the rate at which it progresses.

**Figure 1:**
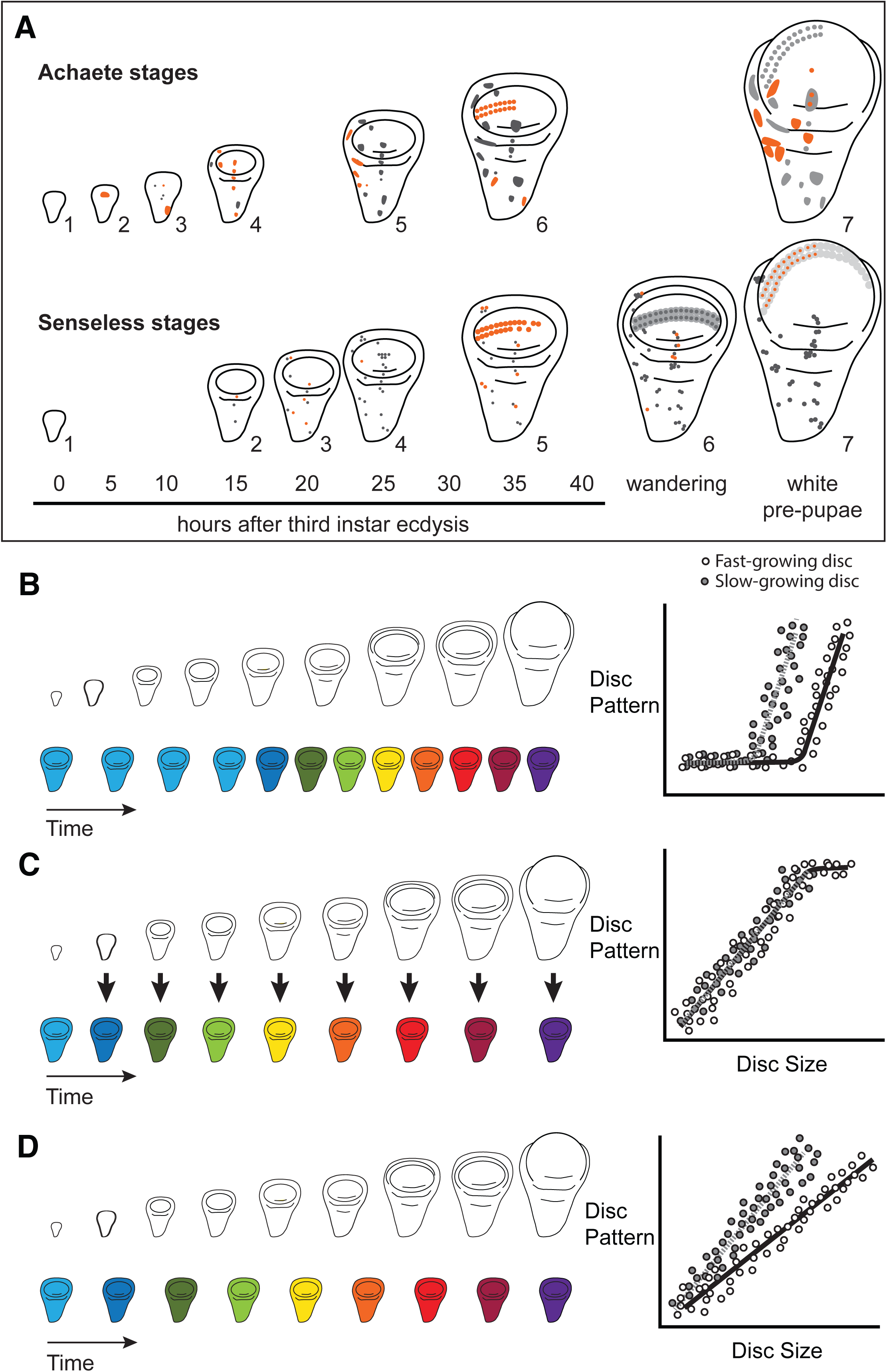
Quantitative assessments of the progression of patterning allow us to test hypotheses about the relationship between the size and patterning stage of the developing wing. A) The staging scheme developed by Oliveira *et al.* 2014 [42] to quantify the progression of Achaete and Senseless pattern. The pattern elements shown in orange are diagnostic for each stage, which is indicated by the number beside the disc. (B-D) The relationship between wing disc size and patterning stage (represented as wing discs progressing through a series of colours) if; (B) Hypothesis 1: Wing discs grow first and then initiate pattern; (C) Hypothesis 2: Wing disc patterning is regulated by wing disc size (arrows); (D) Hypothesis 3: Wing disc pattern and growth are regulated at least partially independently.

The ability to simultaneously quantify both organ growth and pattern allows us to generate, and test, hypotheses regarding how ecdysone coordinates plastic growth with robust pattern. One hypothesis is that growth and patterning occur at different times, with ecdysone driving growth first then pattern later, or vice versa [6]. If this were true, we would expect to identify an interval where ecdysone concentrations primarily affected growth and a second interval where they affected mostly pattern (Figure 1B). There is some precedence for this idea; most of the patterning in the wing discs and ovaries of *D. melanogaster* occurs 15 h after the moult to the third larval instar [60]. Similarly, wing discs are known to grow faster in the early part of the third instar and slow their growth in the mid-to-late third instar [62]. As a second hypothesis, ecdysone could coordinate plastic growth with robust pattern if the impacts of ecdysone on one of these processes depended on its effects on the other. For example, morphogens are known to regulate both growth and patterning of the wing. If ecdysone controlled the action of morphogens, we would expect the progression of patterning would be tightly coupled to growth over time, with different aspects of patterning being initiated at different disc sizes (Figure 1C). Finally, a third hypothesis is that ecdysone regulates growth and patterning of the wing discs independently, and that each process responds in a qualitatively and quantitively different manner to ecdysone [6]. As an example of this, we might see that growth rates increase in a graded response to increasing ecdysone while patterning shows threshold responses, or vice versa. If this were the case, we would expect that growth and the progression of pattern would be uncoupled over time (Figure 1D).

Here we test these hypotheses of whether and how ecdysone co-regulates plastic growth and robust pattern in wing imaginal discs in *D. melanogaster*. We blocked the production of ecdysone by genetically ablating the PG [56], and quantified the effects on growth and patterning rates throughout the third instar. We then manipulated the rate of ecdysone synthesis, by up- or down-regulating the activity of the insulin-signalling pathway in the PG [52, 53], to test how this alters the relationship between disc size and disc pattern. Finally, we tested our hypotheses about how a single steroid can regulate both plastic growth and robust patterning by conducting dose response experiments under two nutritional conditions. These studies provide a foundation for a broader understanding of how developmental hormones coordinate both plastic and robust responses across varying environmental conditions during animal development.

## Results

### Ecdysone is necessary for the progression of growth and patterning

To understand how ecdysone affects the dynamics of growth and patterning, we needed to be able to precisely manipulate ecdysone concentrations. For this reason, we made use of a technique we developed previously to genetically ablate the prothoracic glands (referred to as PGX) [56]. This technique pairs the temperature sensitive repressor of GAL4, GAL80^ts^, with a prothoracic gland-specific GAL4 (*phm-GAL4*) to drive an apoptosis-inducing gene (*UAS-GRIM*). GAL80^ts^ is active at 17°C, where it represses GAL4 action, but inactive above 25°C, which allows *phm-GAL4* to drive expression of *UAS-GRIM* and ablate the PG [63, 64]. Because ecdysone is required at every moult, we reared larvae from egg to third instar at 17°C to repress GAL4, then shifted the larvae to 29°C at the moult to the third instar to generate PGX larvae.

PGX larvae had significantly reduced ecdysteroid titres than control genotypes (Figure 2 Supplement 1). This method of reducing ecdysteroid concentration in the larvae allows us to examine how reducing ecdysone titres affects disc size and pattern in third instar wing imaginal discs, and to manipulate ecdysone concentrations by adding it back in specific concentrations to the food [56]. For simplicity, all the data from the two control strains (either the *phm-GAL4; GAL80ts* or *UAS-GRIM* parental strain crossed to w^1118^), were pooled in all analyses.

**Figure 2:**
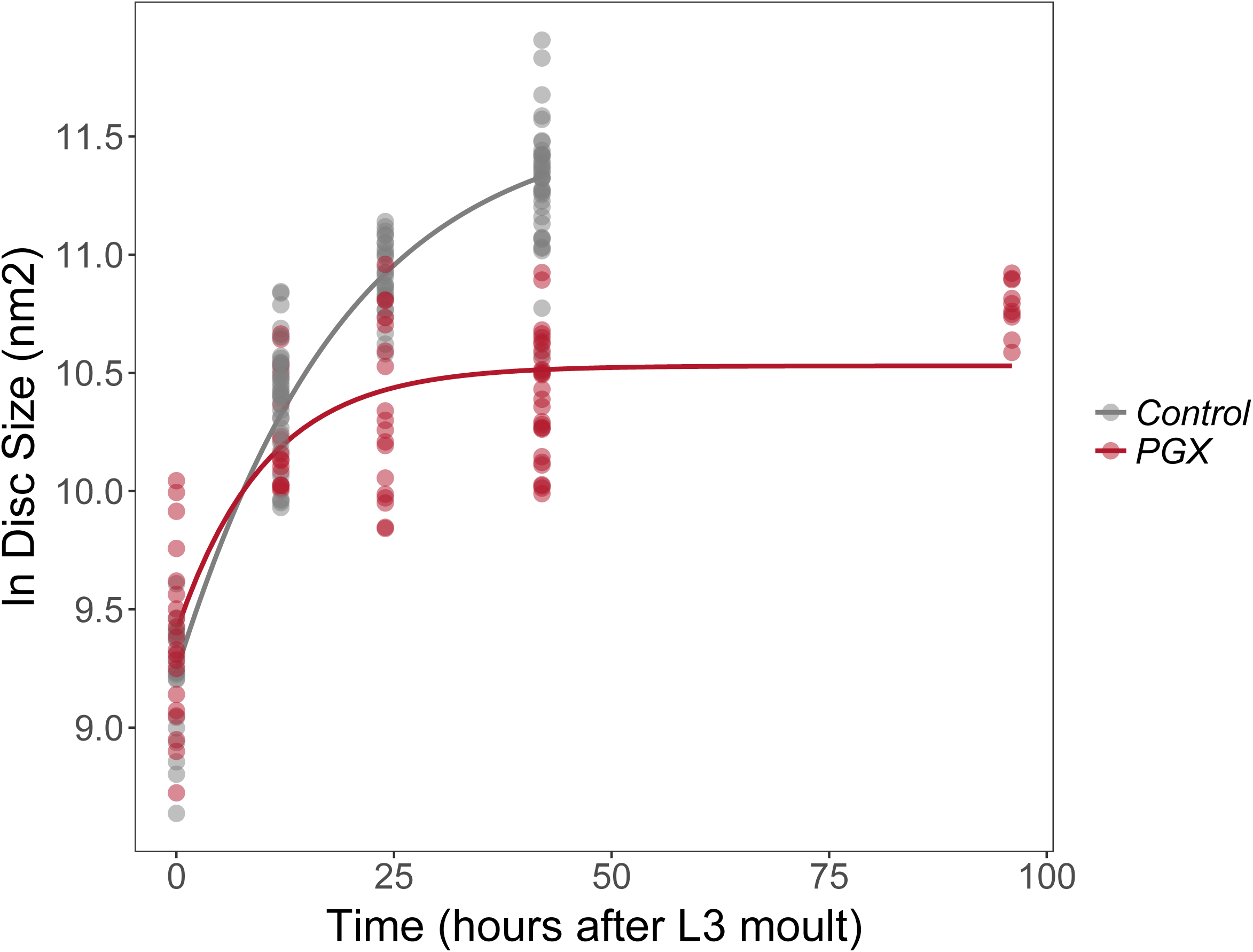
Growth rates of wing discs are reduced in larvae with genetically-ablated PG (PGX) versus control larvae. Curves are Gompertz functions of disc size against disc age (hours after ecdysis to the third larval instar, AEL3). Parameters for the curves are significantly different between PGX and control (Supplementary Table 1). Control genotypes are the pooled results from both parental controls (either the *phm-GAL4; GAL80ts* or *UAS-GRIM* parental strain crossed to w^1118^). Each point represents the size of an individual wing disc. N_PGX_ = 95, N_Control_ = 125 across all time points.

Insect wing discs show damped exponential, or fast-then-slow, growth dynamics [62, 65]. These types of growth dynamics have frequently been modelled using a Gompertz function, which assumes that exponential growth rates slow down with time. The growth of wing discs from control and PGX larvae shows the same pattern, with a Gompertz function providing a significantly better fit to the relationship between log disc size and time than a linear function (ANOVA, linear vs. Gompertz model, *n*> 93, *F*>65, *P*<0.001 for discs from both PGX and control larvae).

Growth of the discs, however, followed a significantly different trajectory in PGX versus control larvae (Figure 2, Supplementary Table 1). In control larvae, discs continue to grow until 42h after ecdysis to the third instar (AEL3), when the larvae pupariate. In contrast, the wing imaginal discs of the PGX larvae grow at slower rates between 0 and 25h AEL3 (Supplementary Table 1) and stop growing at approximately 25h AEL3 at a significantly smaller size. This is despite the fact that PGX larvae do not pupariate, and so disc growth is not truncated by metamorphosis.

We next explored how loss of ecdysone affected the progression of wing patterning. We used the staging scheme that we previously devised in [42] to quantify the progression of wing disc patterning in PGX and control larvae. We selected two genes from this scheme, Achaete and Senseless, as they each progress through seven stages throughout the third instar. Further we can stain for both antigens in the same discs, which allowed us to compare disc size, Achaete stage, and Senseless stage in the same sample.

The progression of Achaete patterning was best fit by a Gompertz function for discs from both PGX and control larvae (ANOVA, linear versus Gompertz model, *n*> 48, *F*>10.4, *P*=0.002) [42], and was significantly affected by reduced ecdysone titres in PGX larvae. In control larvae, the wing discs progressed to Achaete stage 6 or 7 out of 7 stages by 42h AEL3, while in PGX larvae, discs of the same age had not passed Achaete stage 3, and had not matured past Achaete stage 5 by 92h AEL3 (Figure 3A, Supplementary Table 2). The progression of Senseless patterning was best fit by a linear model, but again was significantly affected by reduced ecdysone titres. In control larvae, most discs had progressed to Senseless stage 6 out of seven stages by 42h AEL3, while no disc progressed past Senseless stage 2 by 92h AEL3 (Figure 3B, Supplementary Table 3).

**Figure 3:**
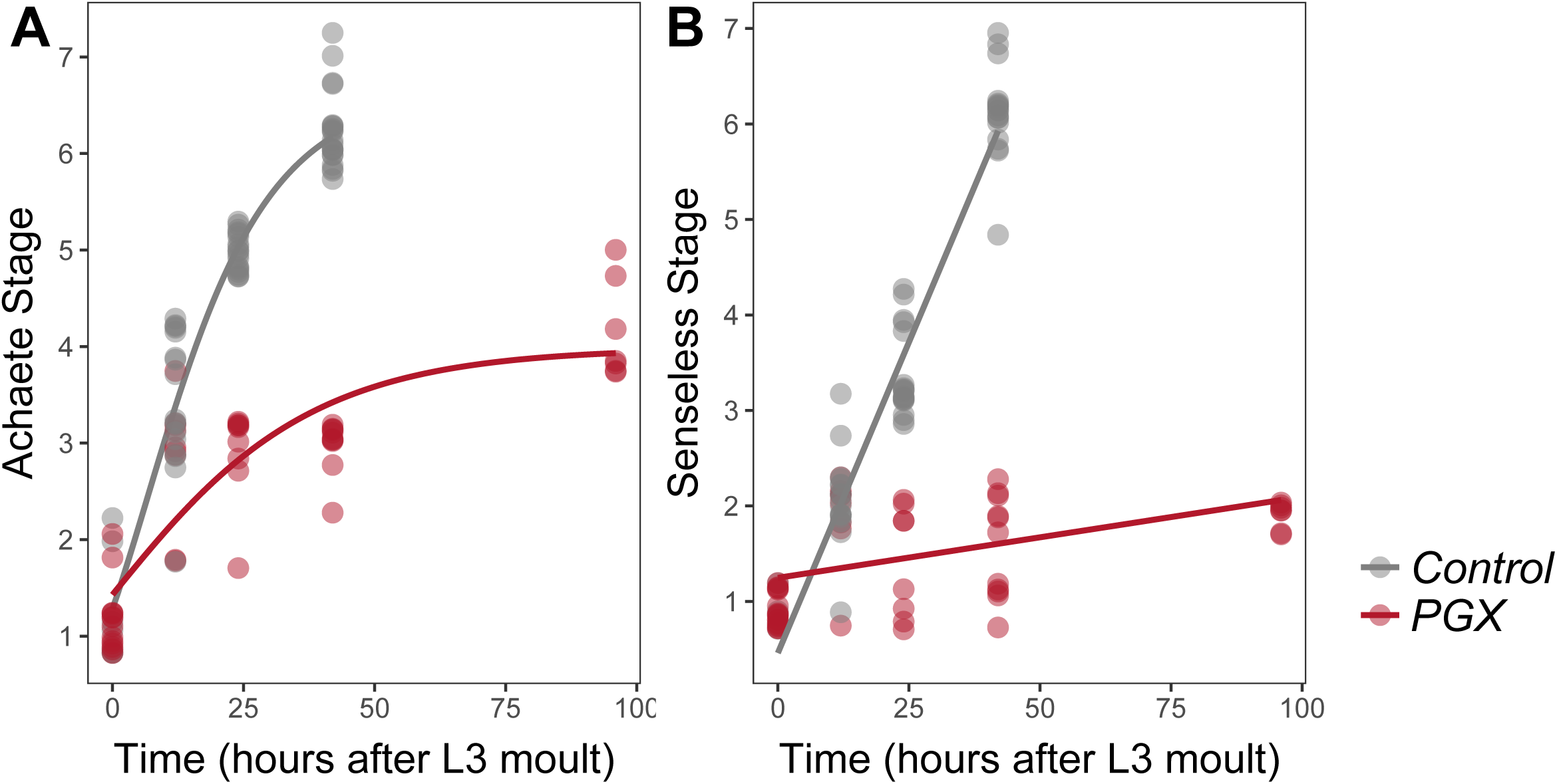
Achaete and Senseless patterning of wing discs is delayed in larvae with genetically-ablated PG (PGX) versus control larvae. (A) Curves are Gompertz functions of Achaete stage against time (hours AEL3). Parameters for the curves are significantly different between PGX and control (Supplementary Table 2). (B) Lines are linear regression of Senseless stage against time (hours AEL3). Parameters for the lines are significantly different between PGX and control (Supplementary Table 3). Control genotypes are the pooled results from both parental controls (either the *phm-GAL4; GAL80ts* or *UAS-GRIM* parental strain crossed to w^1118^). For Achaete: N_PGX_ = 50, N_Control_ = 61, for Senseless: N_PGX_ = 52, N_Control_ = 54 across all time points.

We find no evidence of temporal separation between wing disc growth and the progression of pattern (compare Figures 2 and 3). Both growth and patterning progress at steady rates throughout most of the third instar in control larvae, slowing down only at the later stages of development. Thus, the hypothesis that ecdysone coordinates plastic growth with robust pattern by acting on each process at different times (Figure 1C; Hypothesis 1) is not correct.

To confirm that reduced ecdysone titres were responsible for delayed patterning, and not a systemic response to the death of the glands, we performed a second experiment where we added either the active form of ecdysone, 20-hydroxyecdysone (20E), or ethanol (the carrier) back to the food. PGX and control larvae were transferred onto either 20E or ethanol food and allowed to feed for 42 h, after which we dissected their wing discs and examined their size and pattern. On the control (ethanol) food, wing discs from PGX larvae were smaller (Figure 4A) and showed reduced patterning for both Achaete (Figure 4B) and Senseless (Figure 4C) when compared to control genotypes. Adding 0.15 mg of 20E/mg food fully restored disc size, and Achaete and Senseless pattern, such that they were indistinguishable from control genotypes fed on ecdysone-treated food.

**Figure 4:**
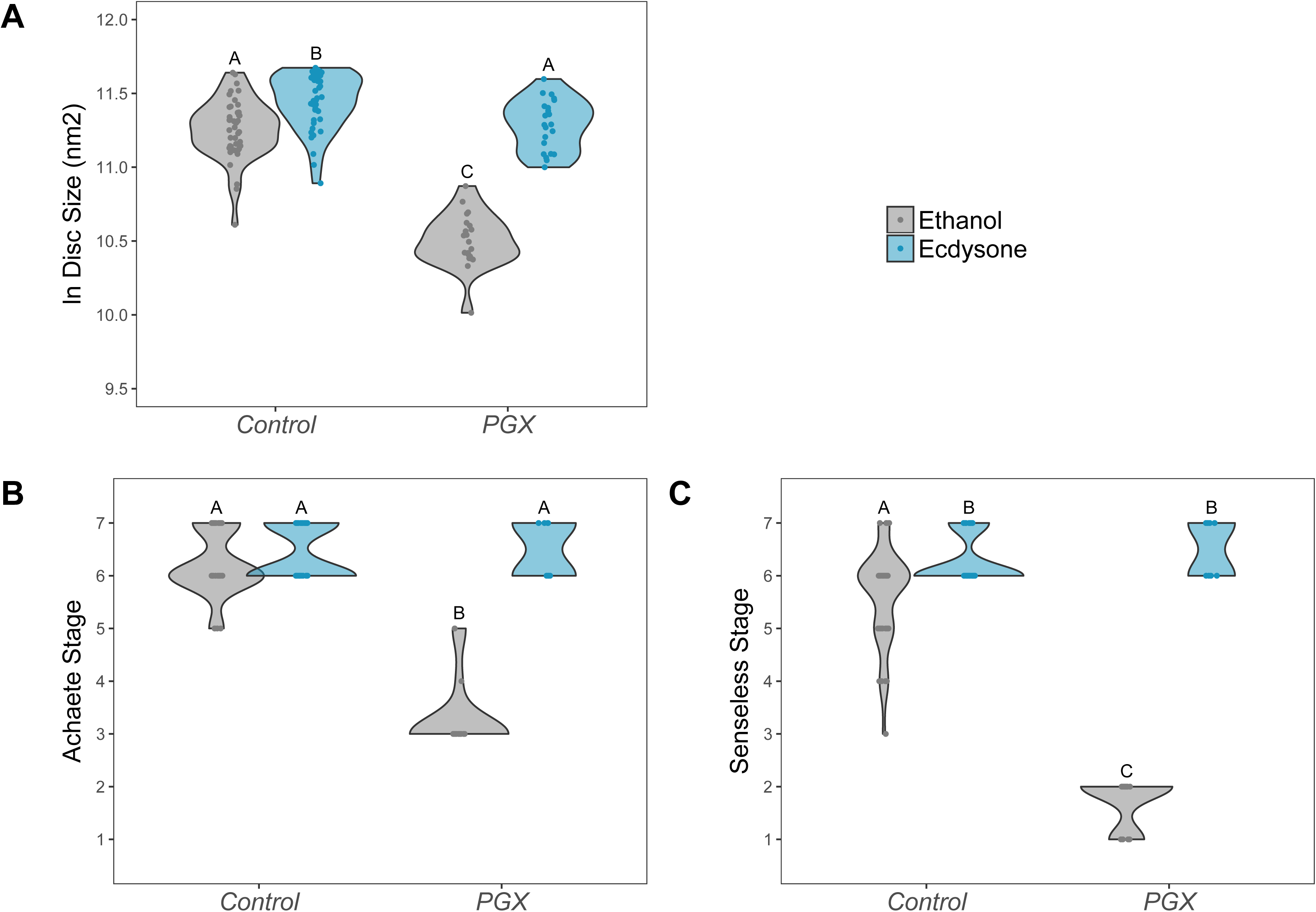
Supplementing PGX larvae with 20-hydroxyecdysone (20E) rescues wing disc growth (A), and Achaete (B) and Senseless (C) patterning. Both the control and PGX larvae were exposed to 20E-treated food (0.15 mg/mg of food) or ethanol-treated food (which contains the same volume of ethanol) at 0 h AL3E. Wing discs were removed at 42 h AL3E. Control genotypes are the pooled results from both parental controls (either the *phm-GAL4; GAL80ts* or *UAS-GRIM* parental strain crossed to w^1118^). Treatments marked with different letters are significantly different (Tukey HSD, *P*< 0.05, for ANOVA see Supplementary Table 4). Data were plotted using violin plots with individual wing discs displayed over the plots. N_PGX + ethanol_ = 21, N_PGX + 20E_ = 23, N_Control + ethanol_ = 43, N_Control + 20E_ = 42.

Collectively, these data indicate that ecdysone is necessary for the normal progression of growth and patterning in wing imaginal discs. The loss of ecdysone has a more potent effect on patterning, however, which is effectively shut-down in PGX larvae, than on disc growth, which continues, albeit at a slower rate, for the first 24 hours of the third instar in PGX larvae.

### Ecdysone rescues patterning and some growth in wing discs of yeast-starved larvae

The observation that ecdysone is necessary to drive both normal growth and patterning suggests that it may play a role in coordinating growth and patterning across environmental conditions. However, to do so it must lie downstream of the physiological mechanisms that sense and respond to environmental change. As discussed above, ecdysone synthesis is regulated by the activity of the insulin-signalling pathway, which is in turn regulated by nutrition. Starving larvae of yeast early in the third instar both suppresses insulin-signalling and inhibits growth and patterning of organs [59, 60]. We explored whether ecdysone was able to rescue some of this inhibition by transferring larvae immediately after the moult to 1% sucrose food that contained either 20E or ethanol and comparing their growth and patterning after 24 hours to wing discs from larvae fed on normal food. Both the PGX and control genotype failed to grow and pattern on the 1% sucrose with ethanol (Figure 5A-C). Adding 20E to the 1% sucrose food rescued Achaete and Senseless patterning in both the control and the PGX larvae to levels seen in fed controls (Figure 5B, C). 20E also partially rescued disc growth in PGX larvae, although not to the levels of the fed controls (Figure 5A). Collectively, these data suggest that the effect of nutrition on growth and patterning is at least partially mediated through ecdysone.

**Figure 5:**
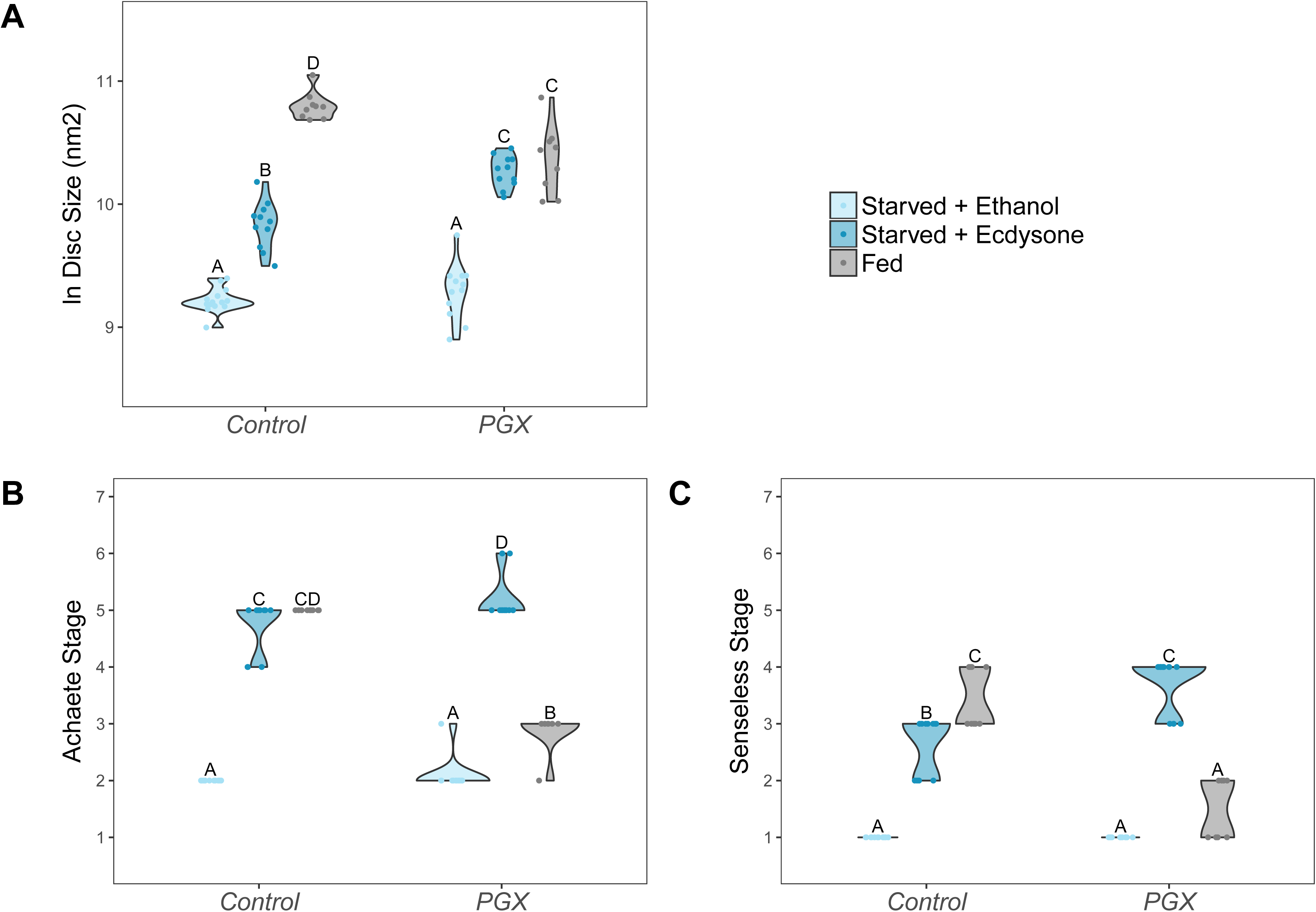
Supplementing PGX larvae with 20-hydroxyecdysone (20E) is able to partially rescue the effect of yeast starvation on (A) wing discs growth, and fully rescue (B) Achaete, and (C) Senseless patterning. Both the control and PGX larvae were exposed from 0 h AL3E to one of three food types: 1) starved + 20E - starvation medium containing 1% sucrose and 1% agar laced with 20E (0.15 mg/mg of food), 2) starved + ethanol - starvation medium treated with the same volume of ethanol, or 3) fed - normal fly food. Wing discs were removed at 24 h AL3E. Control genotypes are the pooled results from both parental controls (the *UAS-GRIM* parental strain crossed to w^1118^). Treatments marked with different letters are significantly different (Tukey HSD, *P*< 0.05, for ANOVA see Supplementary Table 5). Data were plotted using violin plots with individual wing discs displayed over the plots. N_PGX + starved - ethanol_ = 23, N_PGX + starved - 20E_ = 22, N_PGX + fed_ = 26, N_Control + starved-ethanol_ = 28, N_Control + starved-20E_ = 22, N_Control + fed_ = 27.

An important aspect of these data is that in PGX larvae, either supplementing the 1% sucrose food with 20E or feeding them on normal food both rescued wing disc growth (Figure 5A), albeit incompletely. This suggests that nutrition can drive growth through mechanisms independent of ecdysone, and vice versa. In contrast, nutrition alone only marginally promoted Achaete and Senseless patterning in starved PGX larvae, while 20E alone completely restored patterning. Further, even early patterning does not progress in PGX larvae (Figure 3). Thus, the effect of nutrition on patterning appears to be wholly mediated by ecdysone, while the effect of nutrition on growth appears to be partially mediated by ecdysone and partially through another independent mechanism. Ecdysone-independent growth appears to occur early in the third larval instar, however, since disc growth in PGX and control larvae is more-or-less the same in the first 12 hours after ecdysis to L3 (Figure 2).

### Ecdysone drives growth and patterning independently

The data above suggest a model of growth and patterning, where both ecdysone and nutrition can drive growth, but where patterning is driven by ecdysone. We next focused on exploring how ecdysone regulates both growth and patterning. Patterning genes, particularly morphogens, are known to regulate growth, so one hypothesis is that ecdysone promotes patterning, which in turn promotes the ecdysone-driven component of disc growth. A second related hypothesis is that ecdysone-driven growth is necessary to promote patterning. Under either of these hypotheses, because the mechanisms regulating patterning and growth are inter-dependent, we would expect that changes in ecdysone levels would not change the relationship between disc size and disc pattern. An alternative hypothesis, therefore, is that ecdysone promotes growth and patterning through at least partially independent mechanisms. Under this hypothesis the relationship between size and patterning may change at different levels of ecdysone.

To distinguish between these two hypotheses, we increased or decreased the activity of the insulin-signalling pathway in the PG, which is known to increase or decrease the level of circulating ecdysone, respectively [50, 53, 66, 67]. We then looked at how these manipulations affected the relationship between disc size and disc pattern, again focusing on Achaete and Senseless patterning. We increased insulin signalling in the PG by overexpressing InR (*phm>InR*), and reduced insulin signalling by overexpressing the negative regulator of insulin signalling PTEN (*P0206>PTEN*).

We found that a linear model is sufficient to capture the relationship between disc size and Achaete stage when we either increase (*phm>InR*: AIC_linear_ – AIC_logistic_ = 22, ANOVA, *F*_(25,27)_ = 1.71, *P* = 0.2018) or decrease ecdysone synthesis rates (*P0206>PTEN*). Changing ecdysone levels, however, significantly changed the parameters of the linear model and altered the relationship between disc size and Achaete pattern. Specifically, increasing ecdysone level shifted the relationship so that later stages of Achaete patterning occurred in smaller discs (Figure 6A, Supplementary Table 6).

**Figure 6:**
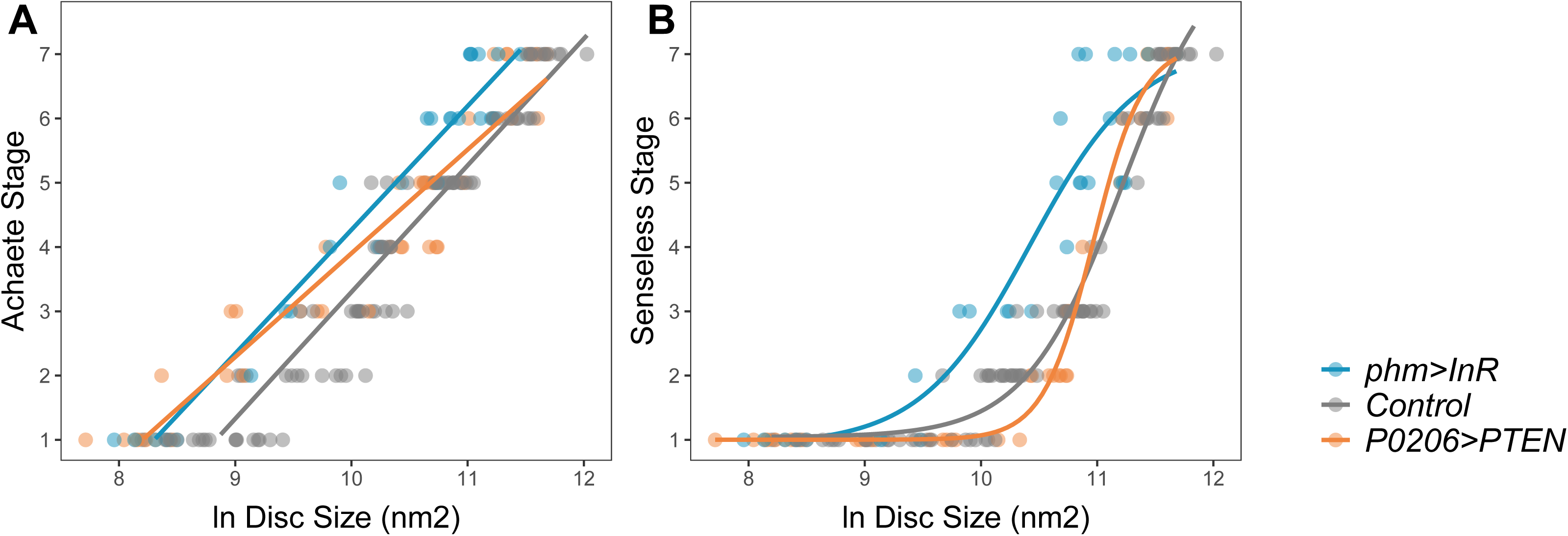
Changing level of ecdysteroidgenesis change the relationship between disc pattern and disc size throughout the third instar larval stage. (A) The relationship between Achaete stage and disc size was fitted with a linear regression, the parameters of which are significantly different between genotypes (Supplementary Table 6). (B) The relationship between Senseless stage and disc size was fitted with a four parameter logistic regression, the parameters of which are significantly different between genotypes (Supplementary Table 7). We staged third instar larvae from the onset of the moult to the formation of white prepupae. The length of this developmental interval varied across genotypes. For control genotypes, we sampled larvae at 0, 10, 20, 30, 48, and 51 (InR control)/53 (PTEN control) h AL3E. For phm>InR, we sampled larvae at 0, 10, 20, 29, 30, and 36 h AL3E. For P0206>PTEN, we sampled larvae at 0, 10, 20, 30, 40, 60, 73, and 80 h AL3E. The number of discs sampled for each genotype and patterning gene was: for Acheate: N_phm>InR_ = 29, N_Control_ = 99, N_P0206>PTEN_ = 40, for Senseless: N_phm>InR_ = 29, N_Control_ = 99, N_P0206>PTEN_ = 41.

The relationship between Senseless pattern and disc size is best fit using a four-parameter logistic (threshold) function, which provides a significantly better fit to the data than a linear function (AIC_linear_ – AIC_logistic_ = 32.2; ANOVA, *F*_(44,46)_ = 25.8, *P* < 0.001). Changing ecdysone levels significantly changed the parameters of the logistic model and altered the relationship between disc size and Senseless pattern (Figure 6B, Supplementary Table 7). Again, increasing ecdysone level shifted the relationship so that later stages of Senseless patterning occurred in smaller discs. Collectively, these data support the hypothesis that ecdysone acts on growth and patterning at least partially independently, and that patterning is not regulated by wing disc size (Figure 1D; Hypothesis 2).

### Ecdysone regulates disc growth and disc patterning through different mechanisms

The data above support a model whereby environmental signals act through ecdysone to co-regulate growth and patterning, generating organs of variable size but invariable pattern. Further, growth is also regulated by an ecdysone independent mechanism, enabling similar progressions of pattern across discs of different sizes. An added nuance, however, is that ecdysone levels are not constant throughout development. Rather, the ecdysone titre fluctuates through a series of peaks throughout the third larval instar and the dynamics of these fluctuations are environmentally sensitive [68]. To gain further insight into how ecdysone co-regulates plasticity and robustness, we therefore explored which aspects of ecdysone dynamics regulate growth and patterning.

Two characteristics of ecdysone fluctuations appear to be important with respect to growth and patterning. First, the timing of the ecdysone peaks set the pace of development, initiating key developmental transitions such as larval wandering and pupariation [53, 67–69]. Second, the basal levels of ecdysone appear to regulate the rate of body growth, with an increase in basal level leading to a reduction in body growth [50, 56, 66, 67, 70]. While several studies, including this one, have established that disc growth is positively regulated by ecdysone [56, 58, 71], whether disc growth is driven by basal levels or peaks of ecdysone is unknown.

There are a number of hypotheses as to how ecdysone levels may drive patterning and growth. One hypothesis is that patterning and ecdysone-regulated disc growth show threshold responses, which are initiated once ecdysone rises above a certain level. This would manifest as low patterning and growth rates when ecdysone titres were sub-threshold, and a sharp, switch-like increase in patterning and growth rates after threshold ecdysone concentrations were reached. Alternatively, both may show a graded response, with patterning and growth rates increasing continuously with increasing ecdysone titres. Finally, disc patterning may show one type of response to ecdysone, while disc growth may show another. Separating these hypotheses requires the ability to titrate levels of ecdysone.

To do this, we reared PGX larvae on standard food supplemented with a range of 20E concentrations (0, 6.25, 12.5, 25, 50 and 100 ng of ecdysone/mg of food). However, as noted above, disc growth early in the third larval instar is only moderately affected by ablation of the PG, potentially obfuscating the effects of supplemental 20E. In contrast, discs from starved PGX larvae show no growth or patterning without supplemental 20E. We therefore also reared PGX larvae on either on standard food or on 20% sucrose / 1% agar medium (from here on referred to as ‘starved’ larvae) supplemented with a range of 20E concentrations. For both control genotypes and PGX larvae, increasing the concentration of 20E in the food increased ecdysteroids titres in the larvae (Figure 7 Supplement 1, Supplementary Table 6).

To quantify the effects of 20E concentration in the on wing disc growth, we dissected discs at 5 h intervals starting immediately after the moult to the third instar (0 h AL3E) to 20 h AL3E. Because male and female larvae show differences in wing disc growth [72], we separated the sexes in this experiment and focused our analysis on female wing discs.

As before, in both PGX and control larvae, wing disc growth was suppressed by starvation (Figure 7 Supplement 2, Supplementary Table 9). To explore how disc size changed over time with increasing 20E concentration, we modelled the data using a second order polynomial regression for time after third instar ecdysis. Increasing the concentration of supplemental 20E increased the disc growth rate in starved PGX larvae (Figure 7A, Supplementary Table 10). In contrast, increasing 20E concentrations had no effect on disc growth rate in fed PGX larvae (Figure 7 Supplement 3, Supplementary Table 10). This confirms that the effect of nutrition on growth hides the effect of 20E early in the third larval instar, and supports that hypothesis that disc growth during this period is primarily regulated by nutrition and only moderately regulated by ecdysone [62].

**Figure 7:**
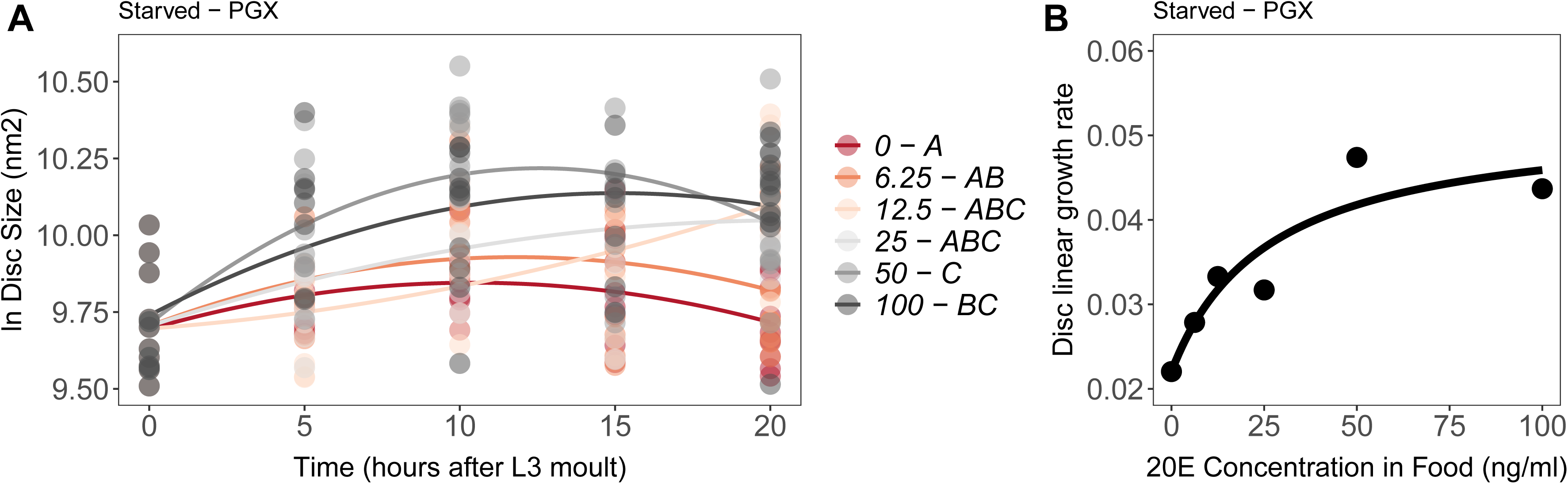
Effect of supplemental 20E on growth of the wing imaginal disc in starved PGX larvae. Growth was modelled as *S* = *E + T + T^2^ + E*T+ E*T^2^*, where *S* = disc size, *E* = 20E concentration, and *T* = disc age. A) There was a significant effect of *E* on the linear growth rate of the wing imaginal discs (Supplementary Table 8 and 9). Each point corresponds to a wing disc, N_PGX – starved_ = 409 (63-72 discs were sampled per treatment across all time points). 20E treatments that do not share a letter (see legend) are significantly different in patterning rates as determined by post-hoc test on the slopes (Supplementary Table 10). B) The linear growth rate of the wing disc was extracted from the growth model for each concentration of 20E, and modelled using a three parameter Michaelis Menten equation: *y = c + (d-c)/(1+(b/x))*, where *c* is *y* at *x=*0*, d* = *y[max]*, and *b* is *x* where *y* is halfway between *c* and *d*. Linear growth rate increases steadily with ecdysone concentration in the food up until 25 ng of 20E/ml of food, after which growth rate increases more slowly with increasing 20E concentration.

To test whether wing disc growth rates show either a graded or threshold response to 20E concentration in starved PGX larvae, we extracted the linear growth rate coefficients from our models. We then modelled the relationship between growth rate and 20E concentration with three non-linear functions: a graded Michaelis Menten function, and with two threshold functions, three and four parameter log-logistic functions. Finally, we tested which model best fit the data using Akaike Information Criteria (AIC) and Bayesian Information Criteria (BIC) for model selection. The model with the lowest AIC and BIC values best fits the data. When wing disc growth rate was modelled with the graded Michaelis Menten function, both the AIC and BIC values were lower than when it was modelled with either threshold function (Supplementary Table 11). This supports the hypothesis that growth rate increases continuously with increasing 20E concentration, with growth rate plateauing after 20E concentrations reach 25 ng/ml (Figure 7B). Thus, disc growth rate appears to show a graded response to 20E level in the absence of nutrition. This is in line with recent findings from the Teleman lab, that show that proliferation and growth in the wing discs increase with increasing 20E concentration in the diet [73].

The effect of 20E concentration on Achaete patterning was qualitatively different to its effect on growth. As before, Achaete patterning did not progress in either starved or fed PGX larvae (Supplementary Figure 4A). In contrast, Achaete patterning did progress in PGX larvae supplemented with 20E. Patterning rates for Achaete did not differ significantly between 0-6.25 ng/ml (fed) and 0-12.5 ng/ml (starved) of 20E (Figure 8 A&C). Above 25 ng/ml of 20E, Achaete patterning occurred at the same rapid rate in both fed and starved PGX larvae (Figure 8 A&C).

**Figure 8:**
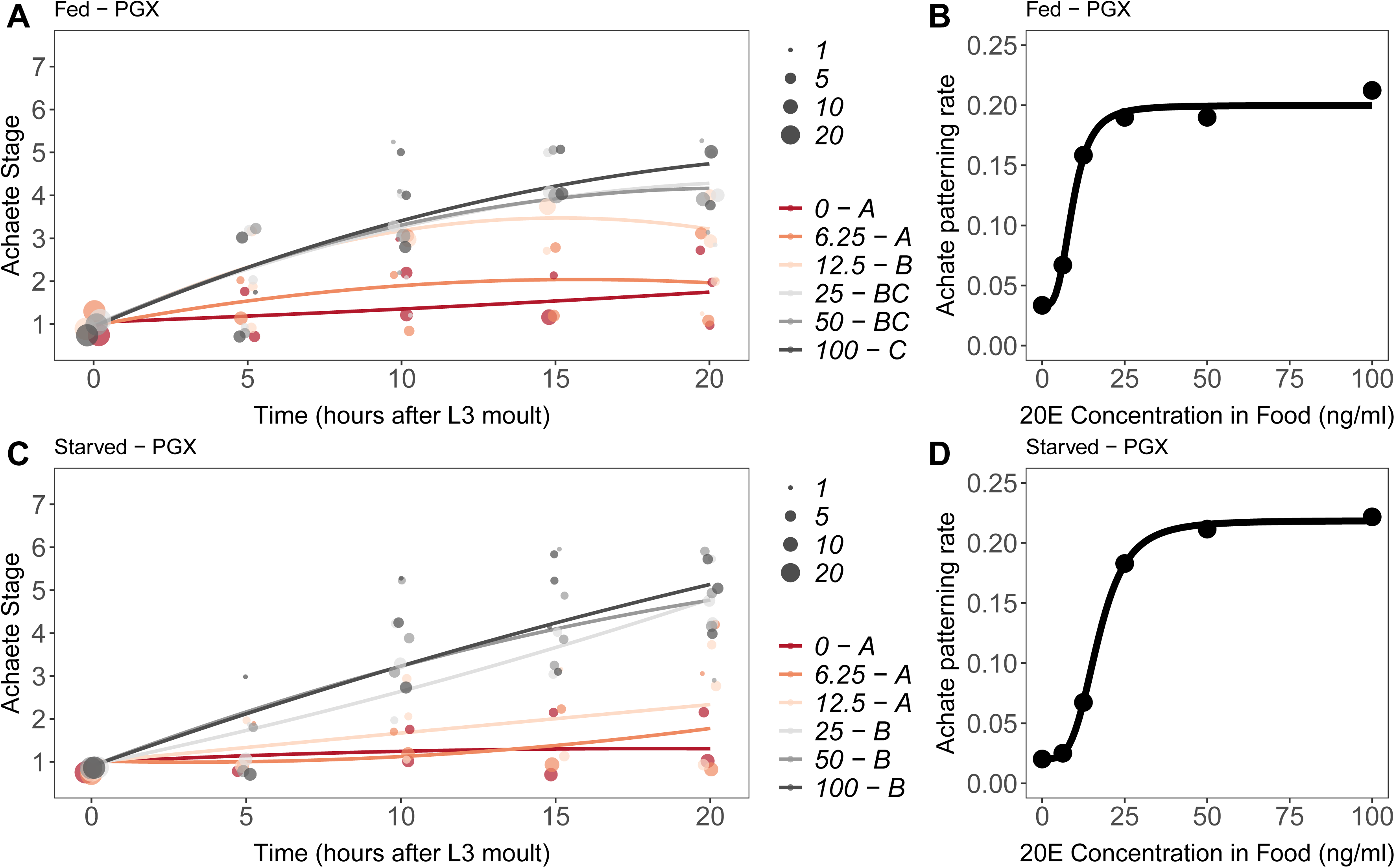
Effect of supplemental 20E on Achaete patterning of the wing imaginal disc in (A-B) fed and (C-D) starved PGX larvae. In A and C, patterning stage was modelled as *S* = *E + T + T^2^ + E*T+ E*T^2^*, where *A* = Achaete stage, *E* = 20E concentration, and *T* = disc age. The size of each point corresponds to the number of wing disc in each category. 20E treatments that do not share a letter (see legend) are significantly different in patterning rates as determined by post-hoc test on the slopes (for ANOVA see Supplementary Table 12). In B and D, we extracted the linear patterning rate from fed (B) or starved (D) PGX larvae. We then modelled the relationship between patterning rate and 20E concentration using a four parameter log-logistic equation: *y = c + (d-c)/(1+e^(b(log(x)-log(a))^)*, where *c* is the lower asymptote*, d* is the upper asymptote, *b* is the rate of increase, and *a* is the inflection point. N_PGX – fed_ = 459, N_PGX – starved_ = 409, 63-86 discs were sampled per treatment across all time points.

To compare the progression of Achaete at different levels of 20E with the progression of disc growth, we modelled the relationship between Achaete pattern, time after third instar ecdysisis, and 20E concentration as second order polynomial regression for fed and starved PGX larvae. As for disc growth, we then extracted the linear coefficients from this model at each level of 20E and modelled the relationship between patterning rate and 20E using a Michaelis Menten function and a three and four parameter log-logistic function. Both the AIC and BIC indicated that that a threshold four parameter log-logistic function fit the data better than a graded Michaelis Menten function (Supplementary Table 9). Thus, unlike growth, Achaete patterning appears to show a threshold response to 20E concentration. Specifically, Achaete patterning is not initiated unless 20E is above a certain level (12.5-25ng/ml), but progresses at the same rate and to the same extent regardless of how high 20E is above this level.

Comparing the timing of Achaete patterning in 20E-supplemented PGX larvae versus fed controls provides some indication of when in normal development the threshold level of 20E necessary to initiate Achaete patterning is reached. Discs from fed control larvae began to reach Achaete stage 4 by 15h AEL3 (Figure 8 Supplement 1B), while discs from both fed and starved PGX larvae supplemented with >25ng/ml of 20E began to reach stage 4 by 10h AEL3 (Figure 8A & C). This suggests that in control larvae, ecdysone levels sufficient to initiate Achaete patterning are only reached 15 h after the mount to the third larval instar.

Senseless patterning did not progress as far as Achaete patterning, only achieving an average of stage 3 in fed and stage 4 in starved larvae at 20 h AL3E when supplemented with 20E. In both fed and starved larvae, supplemental 20E at or below 12.5 ng/ml was insufficient to rescue Senseless patterning, while supplemental 20E at or above 25 ng/ml rescued patterning to approximately the same extent (Figure 9 A&C). As for Achaete patterning, supplemental 20E also initiated Senseless patterning in PGX larvae early when compared to fed controls. Discs from fed control larvae began to reach Senseless stage 3 at 20h AEL3 (Figure 9 Supplement 1), while discs from both fed and starved PGX larvae supplemented with ≥25ng/ml of 20E were at stage 3 by 15h AEL3 (Figure 9A & C).

**Figure 9:**
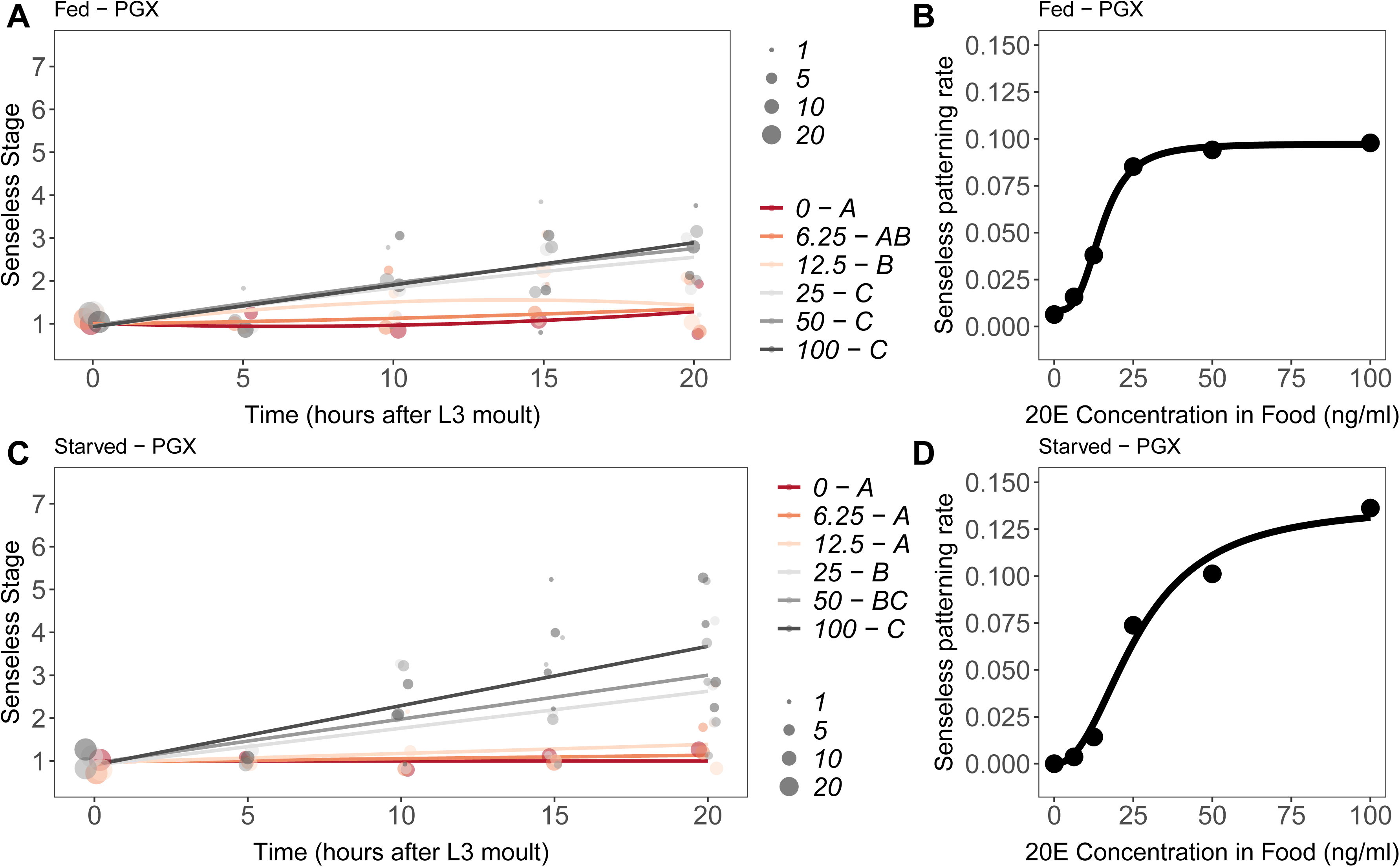
Effect of supplemental 20E on Senseless patterning of the wing imaginal disc in (A-B) fed and (C-D) starved PGX larvae. In A and C, patterning stage was modelled as *A* = *E + T + T^2^ + E*T+ E*T^2^*, where *S* = Senseless stage, *E* = 20E concentration, and *T* = disc age. The size of each point corresponds to the number of wing disc in each category. 20E treatments that do not share a letter (see legend) are significantly different in patterning rates as determined by post-hoc test on the slopes (for ANOVA see Supplementary Table 13). In B and D, we extracted the linear patterning rate in fed (B) or starved (D) larvae. We then modelled the relationship between patterning rate and 20E concentration in B using a four parameter log-logistic equation: *y = c + (d-c)/(1+e^(b(log(x)-log(a))^)*, where *c* is the lower asymptote*, d* is the upper asymptote, *b* is the rate of increase, and *a* is the inflection point. In D, we used a three parameter log-logistic equation: *y = d/(1+e^(b(log(x)-log(a))^)*, where *d* is the upper asymptote, *b* is the rate of increase, and *a* is the inflection point. N_PGX – fed_ = 459, N_PGX – starved_ = 409, 63-86 discs were sampled per treatment across all time points.

We again used a second order polynomial regression to model the relationship between Senseless pattern, time after third instar ecdysis, and 20E concentration. The relationship between the linear coefficient from our model for Senseless patterning and 20E were best fit with a log-logistic rather than a Michaelis Menten function. For fed PGX larvae, the four parameter logistic function provided the best fit to the data (Figure 9B, Supplementary Table 11), whereas for starved PGX larvae, the three parameter logistic function was the best fit (Figure 9D, Supplementary Table 11. Thus, like Achaete patterning rate, Senseless patterning rate shows threshold responses to 20E concentration.

Collectively, these data support a model of disc growth and patterning where ecdysone regulates disc growth as a graded response to basal levels of ecdysone, while ecdysone regulates disc patterning as a single threshold response.

## Discussion

Organs are remarkably good at achieving correct pattern across a broad range of environmental conditions that generate variation in size. While this might seem a simple feat when growth and patterning occur at separate times or are regulated by different hormones, it is considerably less simple if both growth and patterning occur at the same time and are regulated by the same endocrine signal. In this work, we explored how the wing discs of developing *D. melanogaster* use the same hormonal signal to coordinate both their growth and their progression of pattern. We found that ecdysone simultaneously regulates the plastic growth and robust patterning of the wing disc through independent mechanisms: plastic growth responds to ecdysone with a graded response, while robust patterning responds with a single threshold response. We propose that these differences in response represent a potentially general mechanism through which high levels of variation in one organ characteristic, for example size, could be coordinated with low levels of variation in another characteristic of the same organ, for example pattern.

These data make an important contribution to our understanding of how environmental factors, specifically nutrition, affect growth and patterning in developing organs in *Drosophila*. During normal development, circulating ecdysone levels are low during the first 8 hours of the third larval instar, until attainment of a critical size initiates a hormonal cascade that causes ecdysone to fluctuate through a series of characteristic peaks. Each of these peaks is associated with key developmental transitions and prepares the larva for metamorphosis. Low nutrition delays attainment of critical size and the initiation of these peaks, but also appears to raise basal levels of circulating ecdysone between these peaks [74], which slows the growth of the body [74]. At the same time, low nutrition also lowers levels of circulating insulin-like peptides, further slowing growth of the body. While low levels of insulin signalling will suppress imaginal disc growth, the increase in ecdysone concentrations resulting from starvation opposes some of these effects [56, 58, 71] by promoting imaginal disc growth.

Our data suggest that these opposing effects are critical to robust patterning of the wing under different nutritional conditions. At low nutritional conditions, low insulin-signalling at the beginning of the third larval instar slows growth of the body and of the imaginal discs. At this stage, growth of the wing imaginal discs is less dependent on ecdysone [62], evident from the more moderate effects on growth of the wing discs during this period in fed PGX larvae. In the middle of the third larval instar, however, low nutritional conditions elevate basal levels of ecdysone [74]. This drives disc growth independent of insulin-signalling to ensure the discs are of sufficient size to generate viable adult appendages, even as elevated ecdysone suppresses growth of the body as a whole [50, 66, 67]. At the same time, changes in the tempo of the ecdysone fluctuations may ensure that patterning is initiated at the appropriate developmental time, when discs are a sufficient size to generate a viable adult appendage. Three factors therefore appear necessary to achieve variable size but robust patterning under a range of nutritional conditions: (1) a graded growth response to ecdysone; (2) nutritionally-sensitive growth that is independent of ecdysone; and (3) a threshold patterning response to ecdysone.

There is some evidence that our findings apply to patterning and growth of the wings in other insect species. In the tobacco hornworm *Manduca sexta* and the buck-eyed butterfly *Junonia coenia*, wing disc growth is regulated by both ecdysone and insulin (73,76). In the butterfly *J. coenia,* the patterning stage of wing discs can be quantified by the extent of tracheal invasion resulting in wing vein patterning [75]. In this species, wing vein patterning progresses independently of wing size in starved versus fed caterpillars [75]. Thus, the independent regulation of growth and patterning, with growth regulated by both insulin and ecdysone signalling, may be a general mechanism to achieve robust patterning across a range of wing sizes.

While ecdysone and insulin signalling provide systemic cues that tune organ growth to the environmental conditions, morphogens like Wingless and Decapentaplegic (Dpp) act to regulate growth in an organ autonomous manner. The extent to which morphogen gradients respond to these systemic cues is unclear, although the activity of morphogens is known to interact with those of systemic signals at the level of the target genes. For example, insulin/TOR-signalling regulates the activity of Yorkie, a downstream effector of patterning morphogens, including Wingless and Dpp, which controls the rate of cell division [76]. Similarly, reducing ecdysone signalling in the wing reduces the expression of Wingless and reduces Dpp signalling, measured by the levels of phosphorylated Mothers against Dpp expression [56, 58, 59]. Taken together, the signalling pathways that regulate organ growth in response to environmental conditions interact in complex ways with those that regulate organ-autonomous growth suggesting that these two growth regulating mechanisms are not as independent as previously thought [6].

Although the growth of the disc relies on insulin and ecdysone signalling, the progression of patterning for Achaete and Senseless in the wing disc appears to be driven by threshold responses to ecdysone. This is not to say that the progression of patterning does not depend on environmental conditions. Indeed, starvation early in the third instar impedes patterning in both the wing and ovary of *D. melanogaster* [59, 60]. However, rather than resulting from a direct effect of insulin signalling on patterning, the block in the progression of pattern occurs because insulin signalling controls the timing of the first ecdysone pulse in the third larval instar [53, 77]. Our results here confirm that patterning requires suprathreshold concentrations of ecdysone to be initiated. Further, the manner in which ecdysone regulates the progression of patterning ensures that it remains robust against further environmental perturbation. By switching on pattern above threshold ecdysone concentrations, the disc can continue to pattern across a range of environmental conditions, even while growth retains sensitivity to those conditions.

A similar threshold mechanism appears to regulate patterning in the wing discs of other insects. As for *Drosophila,* the earliest stages of wing patterning depend on nutrition in *J. coenia*. If caterpillars are starved before the wing discs begin to pattern, then their discs remain small and their veins unpatterned [75]. In caterpillars starved at later stages after disc patterning has been initiated, the wing discs are small but reach the same vein patterning stage as those of fed control animals. Whether or not the initiation of patterning in *J. coenia* also depends on ecdysone has yet to be determined.

At first glance, the observation that patterning shows a threshold response to ecdysone may not be surprising. In any given cell, patterning is inherently regulated by threshold responses because the expression of the patterning gene product is either on or off in that cell. However, our patterning scheme considers the progression of patterning across the entire field of cells that make up the wing disc. Cells across the wing disc turn on Achaete and Senseless expression at different times, resulting in a continuous progression of pattern with time [42]. Furthermore, like growth, the progression of pattern can vary in rate depending on environmental and hormonal conditions [42]. Consequently, the progression of patterning could, in principle, also show a graded response to ecdysone levels. The observation that once ecdysone concentrations are above threshold, the rate of patterning for Achaete and Senseless is independent of ecdysone provides evidence that the rate of patterning across an entire organ can also show a threshold response: an assumption that, hitherto, has not been tested.

What determines how progression of pattering unfolds through time is unclear. We did not observe discs progressing from stage 1 immediately to stage 7 within a single 5-hour time interval even at the highest 20E concentrations. This suggests that there are additional temporal factors that regulate the order of patterning progression. Almost certainly, interactions between the gene regulatory networks that regulate patterning control how patterning progresses across regions of the wing disc. We have very little understanding if/how the different regions of the wing communicate with each other to achieve this. In principle, differences between when cells turn on Achaete and Senseless across the disc could arise in response to other developmental signals, such as from the Dpp, Wingless, or Hedgehog morphogen gradients responsible for correctly scaling and patterning the wing.

Part of this temporal signature might arise from ecdysone itself. In this study we exposed animals to tonic concentrations of ecdysone. Developing larvae, however, secrete four pulses of ecdysteroids between the moult to third instar and pupariation [68]. We have little understanding of how developmental information is encoded within these pulses. In principle, individual pulses could either prime tissues to become responsive to hormones, or could alter their sensitivity – as the early ecdysone pulse does for wing disc growth and patterning [59, 60, 62]. Future studies comparing the difference between tonic and phasic exposure to hormone would help clarify the roles of the ecdysone pulses.

While our study has focussed on contrasting the robustness of patterning with plasticity of growth, depending on what is being measured there are instances where we expect patterning to also show plasticity [78]. For example, although the specification of cell types in the correct location within an organ may show little variation across environmental conditions, the number of structures specified can vary. The total number of abdominal and sternopleural bristles varies with temperature [79, 80], as does the number of terminal filament stacks that are specified in the ovary, which is also affected by nutrition [81–84]. Plasticity in the number of bristle cells or terminal filament stacks presumably occurs because the mechanisms that specify the number of each structure do not scale with organ size. In other cases, the location of specific cell types may also be plastic. For example, there is an extensive literature exploring how the relative positions of veins in the wings of *D. melanogaster* and other insects is affected by environmental factors such as nutrition and temperature (e.g. [85–88]). Plasticity in wing shape is likely to be more complex, and may involve process that act at many different points during wing development [89, 90]. Future studies targeting how the mechanisms that establish the position of cell types differ from those that determine the number of cells of a given type would allow us to further define what makes traits either sensitive or robust towards changes in environmental conditions, and at what level.

## Materials and Methods

### Fly stocks and rearing conditions

We initially manipulated growth rates and developmental timing by either altering the rearing temperature or the rates of ecdysone synthesis in developing *D. melanogaster* larvae. We used the isogenic strain, Samarkand (SAM), which also bears the w^1118^ mutation, when rearing larvae under one of three thermal conditions: 18°C, 25 °C, and 29°C. To accelerate the rates of ecdysone synthesis, we used the progeny from *w^1118^*;*phantom-*GAL4, which is expressed in the prothoracic glands, crossed with yw *flp; UAS InR29.4* (*phm > InR*). We decreased rates of ecdysone synthesis by crossing *P0206-GAL4*, which drives expression throughout the ring gland, with *yw; UAS PTEN* (*P0206 > PTEN*). Even though *P0206*-GAL4 is a weaker GAL4 driver for the prothoracic gland and also drives expression in the corpora allata, we chose to use it to drive *UAS PTEN* because *phm>PTEN* larvae die as first instar larvae [52]. The parental lines *yw flp; UAS InR29.4* (+ *> InR*) and *yw; UAS PTEN* (+ *> PTEN*) were used as a reference for the *phm > InR* and *P0206 > PTEN* genotypes respectively.

Flies of the above genotypes were raised from timed egg collections (2-6 hours) on cornmeal/molasses medium containing 45 g of molasses, 75 g of sucrose, 70 g of cornmeal, 10 g of agar, 1100 ml of water, and 25 ml of a 10% Nipagin solution per liter. Larvae were reared at low density (200 eggs per 60 x 15 mm Petri dish) in a 12 h light-dark cycle with 70% humidity, and maintained at 25°C unless stated otherwise.

We used a transgenic combination that allowed us to genetically ablate the prothoracic gland and eliminate native ecdysone synthesis specifically in the third larval instar. We crossed a *tub-GAL80^ts^, phantom GAL4* strain with *UAS Grim* to generate *PGX* progeny [56]. GAL80^ts^ is a repressor of GAL4 active at temperatures lower than 22°C [63]. Rearing PGX larvae at 17°C allows GAL80^ts^ to remain active, thus the *phantom GAL4* cannot drive the expression of *UAS grim* to promote cell death. Under these conditions, larvae can moult, pupariate, and complete metamorphosis [56]. Changing the larval rearing temperature to 29°C disables GAL80^ts^ activity, thus ablating the prothoracic gland [56]. The progeny of the isogenic control strain, *w1118*, crossed with one of two parental lines, either *phantom*-GAL4 (*PG* > +) or *UAS Grim* (+ > *Grim*), were used as controls for genetic background effects. The parental controls were reared under the same thermal conditions as PGX larvae.

Crosses, egg collections, and larval rearing were done on the cornmeal/molasses medium (above) for the experiments in Figures 2-6 or, for the experiments in Figures 7-9, on Sugar-Yeast-Agar (SYA) medium: 50g of autolysed Brewer’s yeast powder (MP Biomedicals), 100 g of sugar, 10 g of agar, and 1200 ml of water. In addition, we added 3 ml of proprionic acid and 3 grams of nipagen to the SYA medium to prevent bacterial and fungal growth. Egg collections were performed on SYA medium for 4 hours at 25°C or overnight at 17°C and larvae were reared at controlled densities of 200 eggs per food plate (60 x 15 mm Petri dish filled with SYA medium) at 17°C, as described previously [56].

### Animal staging and developmental time

To measure the effects of larval rearing temperature or changes in the rates in ecdysone synthesis on wing disc growth and wing disc patterning, larvae were staged into 1-hour cohorts at ecdysis to the third larval instar as in [52, 59]. To do this, food plates were flooded with 20% sucrose and all second instar larvae were transferred to a new food plate. After one hour, the food plate was flooded once again with 20% sucrose and the newly moulted third instar larvae were collected and transferred to new food plates and left to grow until the desired time interval. Animals were staged and their wing discs dissected at defined intervals after the larval moult as in [42].

For the experiments in Figures 7-9, PGX, *phm>,* and *>Grim* genotypes, larvae were raised from egg to second instar at 17°C. Larvae were staged into 2-hour cohorts at ecdysis to the third larval instar using the methods described above. We separated female and male larvae by examining them for the presence of testes, which are significantly larger and more visible even in newly moulted males.

### Exogenous ecdysone feeding treatments

To show that ecdysone could rescue patterning and growth in PGX larvae (Fig 4C&D), we added either 0.15 mg of 20E (Cayman Chemical, Item No. 16145) dissolved in ethanol, or an equivalent volume of ethanol, to 1 ml of standard food. Both the ethanol- and ecdysone-supplemented food were allowed to sit at room temperature for at least 4 hours to evaporate off excess ethanol before use. Twelve larvae were transferred to one of the two supplemented foods either at 0 h AL3E and left to feed for 42 h or at 42 h AL3E and left to feed for 24 h.

To determine the relative contributions of nutrition-dependent signalling or ecdysone to growth and patterning, we fed newly moulted PGX and control larvae 1 ml of starvation medium (1% sucrose with 1% agar) supplemented with either 0.15 mg of 20E dissolved in ethanol, or an equivalent volume of ethanol (Fig Suppl 4). Supplemented food was left at room temperature for at least 4 hours to evaporate excess off ethanol before use. Larvae were collected at 24 h AL3E for tissue dissection.

For the 20E dose response experiments, we conducted an initial pilot that showed that supplementing the food with 100 ng of ecdysone/mg food could rescue most of the Achaete and Senseless patterning in PGX wing discs. We collected newly moulted third instar larvae, separated the sexes, and then transferred 10-20 larvae to either sucrose food (20% sucrose, 1% agar; starved) or SYA food (fed) at 29°C. We fed these larvae on one of six ecdysone concentrations: 0, 6.25, 12.5, 25, 50, or 100 ng of 20E/ mg food. We added the same volume of ethanol to all treatments.

To quantify the relationship between the concentration of 20E administered and the concentration of ecdysteroids in the hemolymph, we allowed newly-ecdysed larvae to feed on either sucrose or SYA food that had been supplemented with one of the six concentrations of ecdysone for 20h at 29°C. We then transferred them onto either sucrose food or SYA food that did not contain ecdysone but was dyed blue. They were left to feed for 2 hours until their guts were filled with blue food. This extra step was taken so that we could be sure that our hemolymph ecdysone titres were not contaminated with ecdysone from the food. Thirty – forty larvae were then weighed as a group, and transferred to 5 times their weight in volume of ice cold methanol. Larvae were homogenized and ecdysone titres were determined using a 20-Hydroxyecdysone Enzyme ImmunoAssay Kit (Cayman Chemical, Item Number 501390) as per manufacturer’s instructions.

### Dissections and immunocytochemistry

For each sample, 10-20 larvae were dissected on ice cold phosphate-buffered saline (PBS) and fixed in 4% formaldehyde in PBS overnight at 4°C. After fixation, the tissue was washed four times (15 minutes per wash) with 0.3% Triton X-100 in PBS (PBT), then blocked for 30 minutes at room temperature in 2% heat-inactivated normal donkey serum in PBT. After blocking, the tissue was incubated in a primary antibody solution diluted with 2% heat-inactivated normal donkey serum in PBT overnight at 4°C. We used the guinea pig anti-Senseless (Nolo et al., 2000, 1:1000) and mouse anti-Achaete (supernatant, 1:10) primary antibodies. To compare signal across tissues, we stained for both antigens simultaneously. The washing and blocking procedure was repeated after primary antibody incubation and then the tissue was incubated in a secondary antibody (1:200 each of anti-guinea pig (546nm) and anti-mouse (488nm)) overnight at 4°C. The tissues were washed with PBT and rinsed with PBS and then the wing imaginal discs were mounted on poly-L-lysine-coated coverslips using Fluoromount-G (SouthernBiotech). Tissues were imaged using either a Leica LSM 510 or a Nikon C1 upright confocal microscope and processed using ImageJ (version 2.0) and Adobe Photoshop CC 2017.

### Quantifications of wing imaginal disc size and Achaete and Senseless pattern

We quantified wing disc size using disc area as a proxy. All quantifications were done using ImageJ. Wing discs show exponential growth in the third instar. Thus, we studied the growth trajectories of the discs by ln-transforming disc area.

Achaete and Senseless stage was quantified using the staging scheme developed by [42], associating each of the wing imaginal discs to an Achaete or Senseless stage varying from 1 to 7.

### Statistical analysis

All the analyses were conducted in R and the annotated R markdown scripts and data for the analyses are deposited on Figshare (DOI: 10.26180/13393676).

For the relationship between time after third instar ecdysis and disc size (log µm^2^) or disc pattern (Achaete or Senseless) we fit either linear or Gompertz models and selected the model that best fit the data, using ANOVA and AIC. The Gompertz model was parameterized as 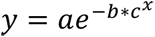, where *y* is disc size/pattern, *x* is time, *a* is the asymptote of *y*, *b* controls where along the x-axis the curve is positioned and *c* is the scaling constant, such that *c* = *e^-g^* where *g* is the growth/patterning rate (thus, the higher *g* the lower *c*). To compare the parameters of linear models between treatments and genotypes we used ANOVA. To compare the parameters of Gompertz models between treatments and genotypes we used ANOVA to compare the fit of models that assign the same constants across groups versus models that assigned group-specific constants.

For the relationship between disc size (log µm^2^) and Senseless pattern we fit a four parameter logistic model parametrized as 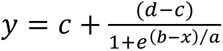 where *y* is disc pattern, *x* is disc size, c is the minimum asymptote, *d* is the maximum asymptote, *b* is the inflection point and *a* is the scaling constant, such that *a* = 1/*k*, where *k* is the logistic growth rate. We again used ANOVA to compare the fit of models that assign the same parameters across groups versus models that assigned group-specific parameters. The relationship between disc size and Achaete pattern was fit using a linear model and compared across treatments using ANOVA.

We used ANOVA to compare disc size/patten at specific time points between treatments and genotypes, using a Tukey’s HSD test to allow comparison among groups.

Finally, to compare the effects of 20E supplementation in the diet on the progression of wing disc growth, Achaete patterning, and Senseless patterning, we fit a second-order orthogonal polynomial regression using disc size/patterning stage as our dependent variable, and 20E concentration and linear and quadratic terms for time as fixed effects. Fitting a single model to the data allowed us to compare the same model parameters for growth and patterning. We then extracted the linear rate of change at each 20E concentration using the *emtrends* function of the *emmeans* package in *R* [91]. The changes in growth/patterning rate with 20E concentration were modelled using three non-linear functions: (1) a continuous Michaelis Menten function: 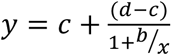, where *c* is *y* at *x=*0*, d* is the maximum asymptote, and *b* is *x* where *y* is halfway between *c* and *d*, (2) a threshold three-parameter log-logistic function: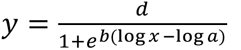 where *d* is the maximum asymptote, *b* is the rate of increase, and *a* is the inflection point, and (3) a threshold four-parameter log-logistic function: 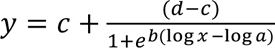 where *c* is the minimum asymptote, *d* is the maximum asymptote, *b* is the rate of increase, and *a* is the inflection point. For each model we calculated the Akaike Information Criteria (AIC) and Bayesian Information Criteria (BIC) to allow model selection. The model that produces the lowest AIC and BIC value best fits the data.

For all parametric tests we checked for homoscedasticity and normality of errors.

## Acknowledgements

We would like to thank members of the Mirth and Shingleton labs, past and present, for their useful discussions relating to this project. This research was supported by NSF grants IOS-0919855, IOS-1557638 and IOS-1952385 to AWS and an ARC Future Fellowship (FT170100259) to CKM.

---

**Figure 2 Supplement 1:**
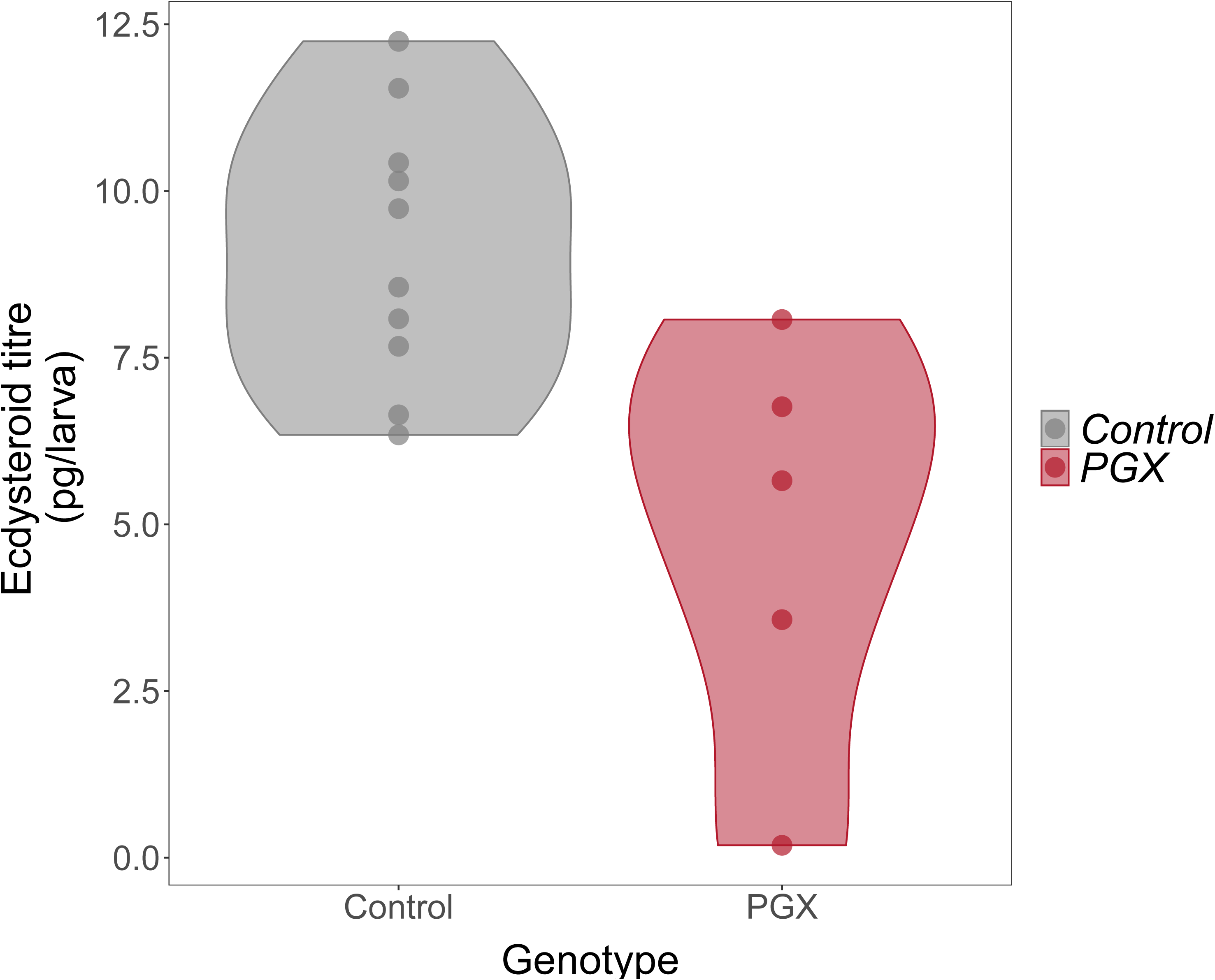
Ecdysteroid titres in PGX and control larvae. Newly ecdysed larvae were placed on sucrose/yeast diets. Ecdysteroid titres in control (phm > + and + > grim) are significantly higher ecdysteroid titres than PGX larvae, as determined by linear models and pairwise comparisons of the means (F value_1, 13_ = 10.75, p value = 0.006, N_control_ = 10, N_PGX_ = 5). Data is plotted with violin plots and the individual replicates (5 per genotype) are included as points overlaying the violin plots.

**Figure 7 Supplement 1:**
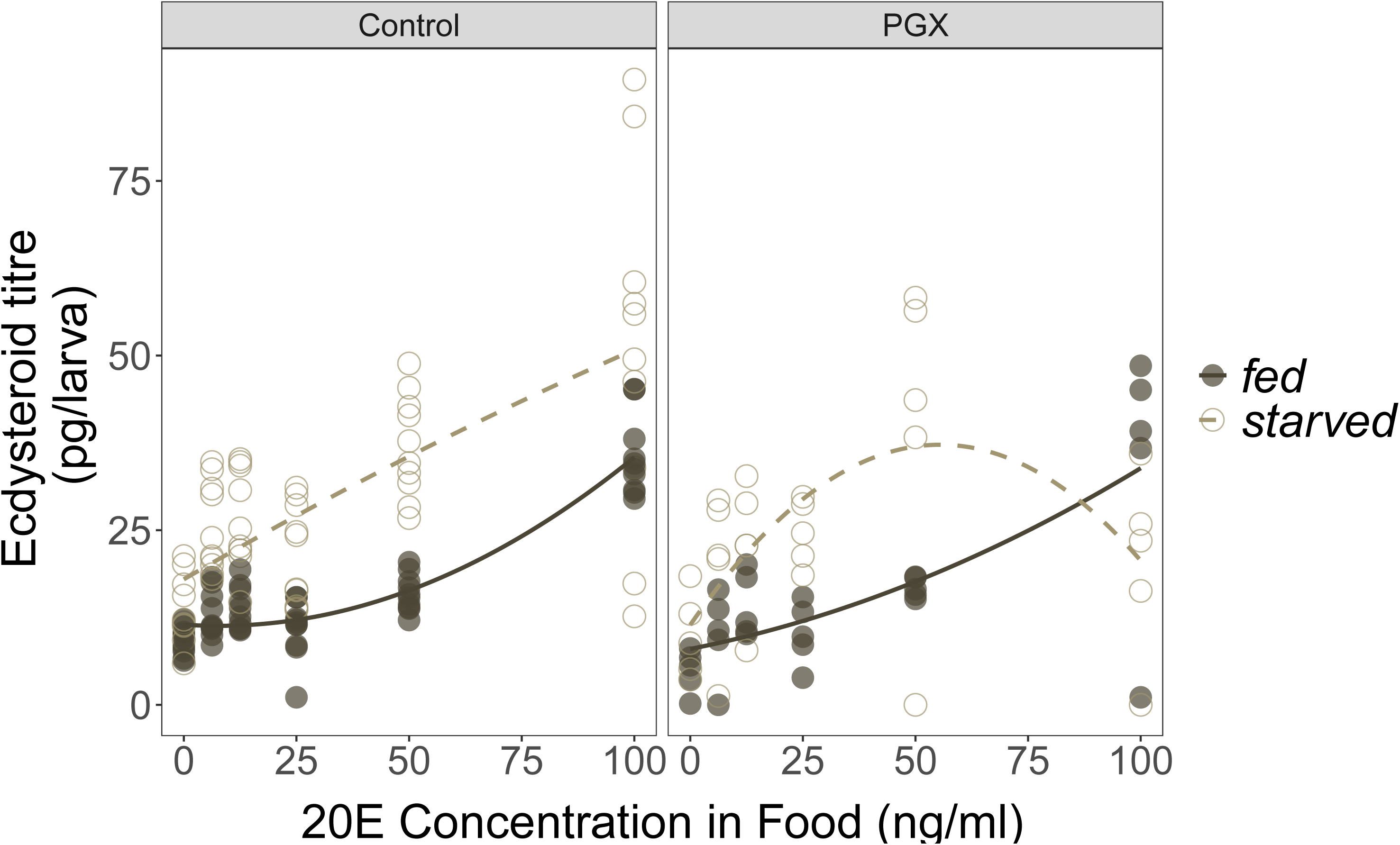
The effects of 20E concentration in the food on ecdysteroid titres in control and PGX larvae. Newly ecdysed larvae were placed on sucrose/yeast (fed)or 20% sucrose/1% agar (starved) diets supplemented with a range of 20E concentrations for 20h. They were then transferred to the same diet without 20E and dyed with blue food colouring for 2 h, to eliminate residual 20E in the gut. Ecdysteroid titres in fed and starved control (phm > + and + > grim) and PGX larvae fed a range of 20E concentrations. There is a significant positive relationship between 20E concentration in the food and the concentration of ecdysteroids in the larvae, as indicated by a significant 20E term (Supplementary Table 8). Furthermore, starved larvae had higher ecdysone titres than fed larvae. Open and closed points represent the biological replicates, and solid and dashed lines are linear regressions. N_control - starved_ = 60, N_control - fed_ = 60, N_PGX - starved_ = 30, N_PGX - fed_ = 30 across all 20E concentrations.

**Figure 7 Supplement 2:**
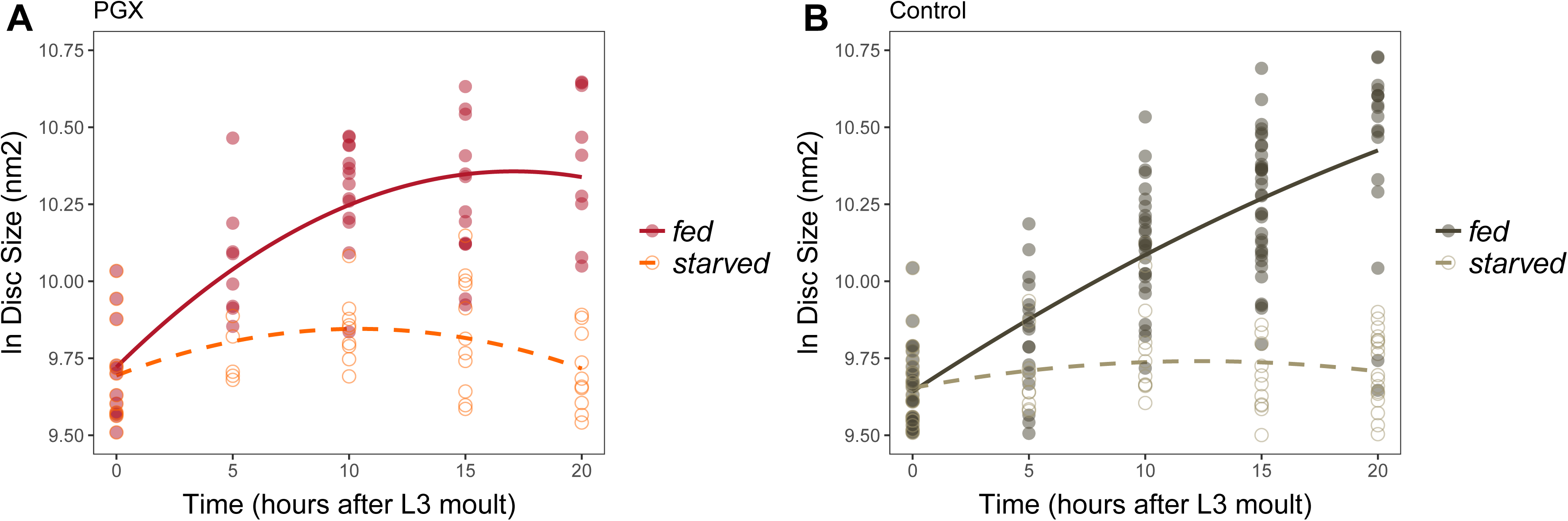
Wing imaginal disc growth is suppressed in fed PGX larvae relative to controls, and in starved larvae of both genotypes. Wing disc growth was modelled as a quadratic, and there was a significant interaction between genotype (PGX v. Control) and nutrition (fed v. starved) on growth (Supplementary Table 9). Solid line/closed point = fed larvae, broken line/open point = starved larvae. Each point corresponds to a wing disc, N_PGX - starved_ = 67, N_PGX - fed_ = 74, N_control - starved_ = 118, N_control - fed_ = 151 across all time points.

**Figure 7 Supplement 3:**
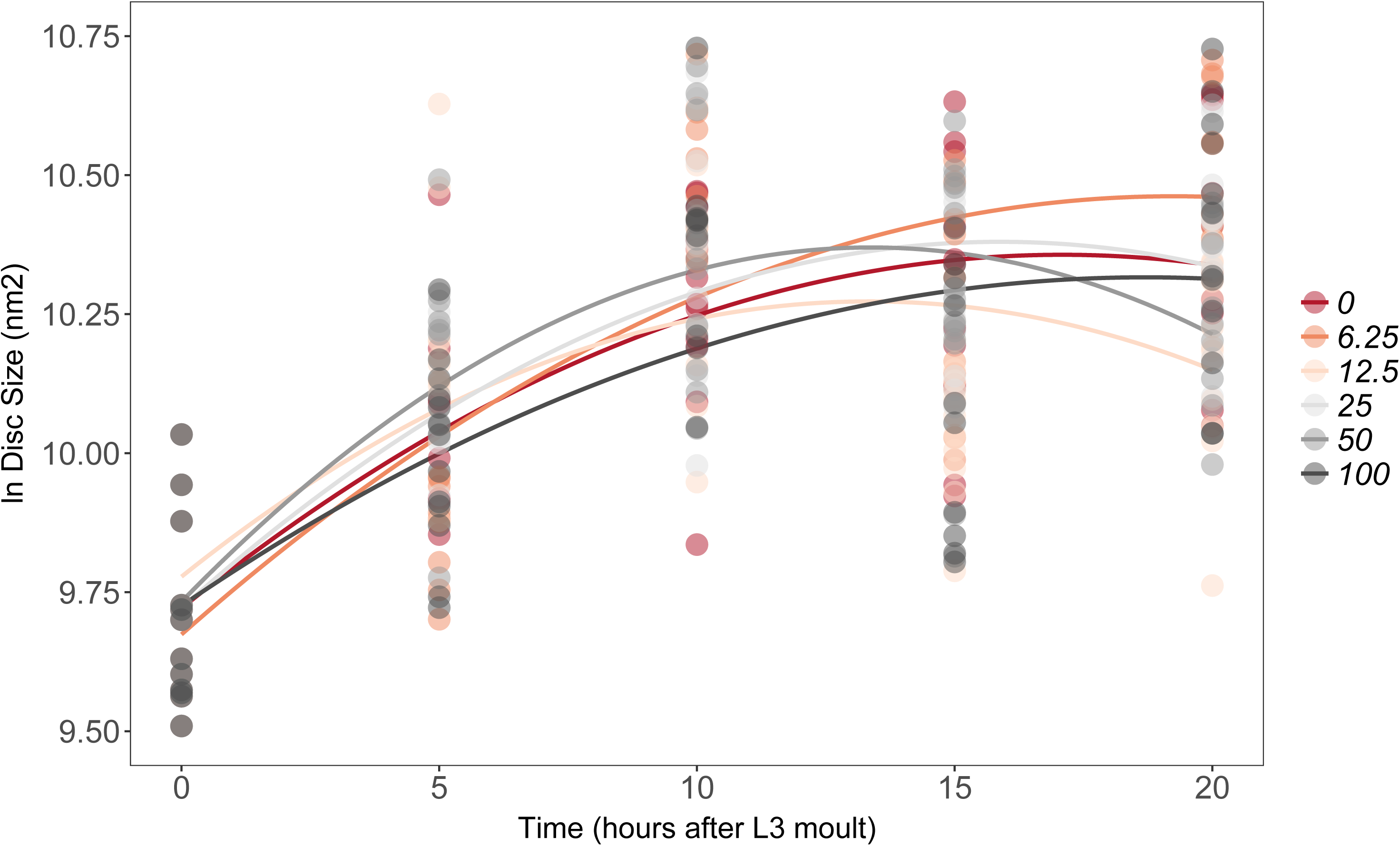
There is no effect of supplemental 20E on growth of the wing imaginal disc in fed PGX larvae. Growth was modelled as *S* = *E + T + T^2^ + E*T+ E*T^2^*, where *S* = disc size, *E* = 20E concentration, and *T* = disc age. There was no significant effect of *E* on the linear or quadratic growth rate of the wing imaginal discs (Supplementary Table 10). Each point corresponds to a wing disc, N_PGX – fed_ = 459 (73-86 discs were sampled per treatment across all time points).

**Figure 8 Supplement 1:**
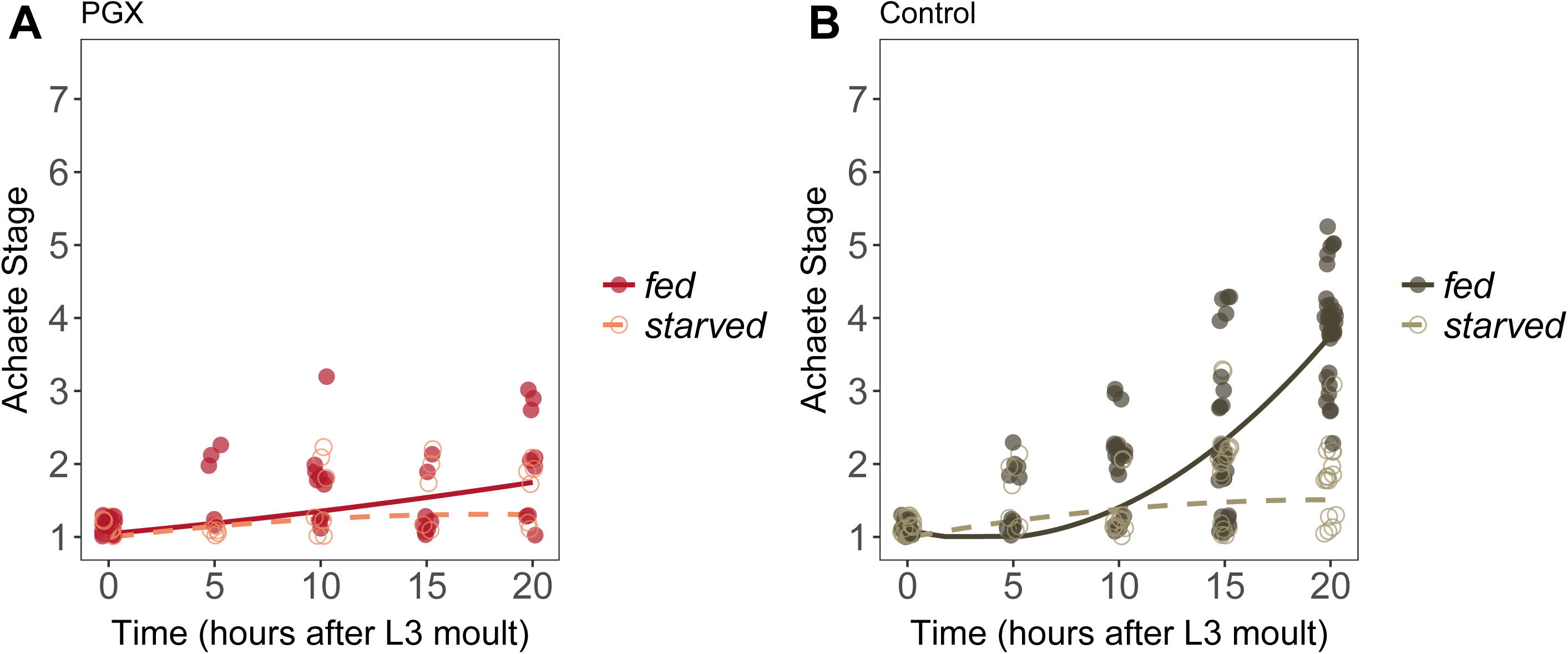
Achaete patterning in wing discs from fed and starved PGX and control larvae. (A) Patterning does not progress in either fed or starved PGX larvae. (B) Patterning does not progress in starved control larvae but does in fed control larvae. There is a significant interaction between the effects of disc age and food on Achaete patterning in control larvae (orthogonal polynomial regression: F_food*disc age_^2^ =67.98, P < 0.001), but not in PGX larvae (orthogonal polynomial regression: F_food*disc age_^2^ =1.81, P = 0.163). Each point corresponds to a wing disc, N_PGX - starved_ = 67, N_PGX - fed_ = 74, N_control - starved_ = 118, N_control - fed_ = 151 across all time points.

**Figure 9 Supplement 1:**
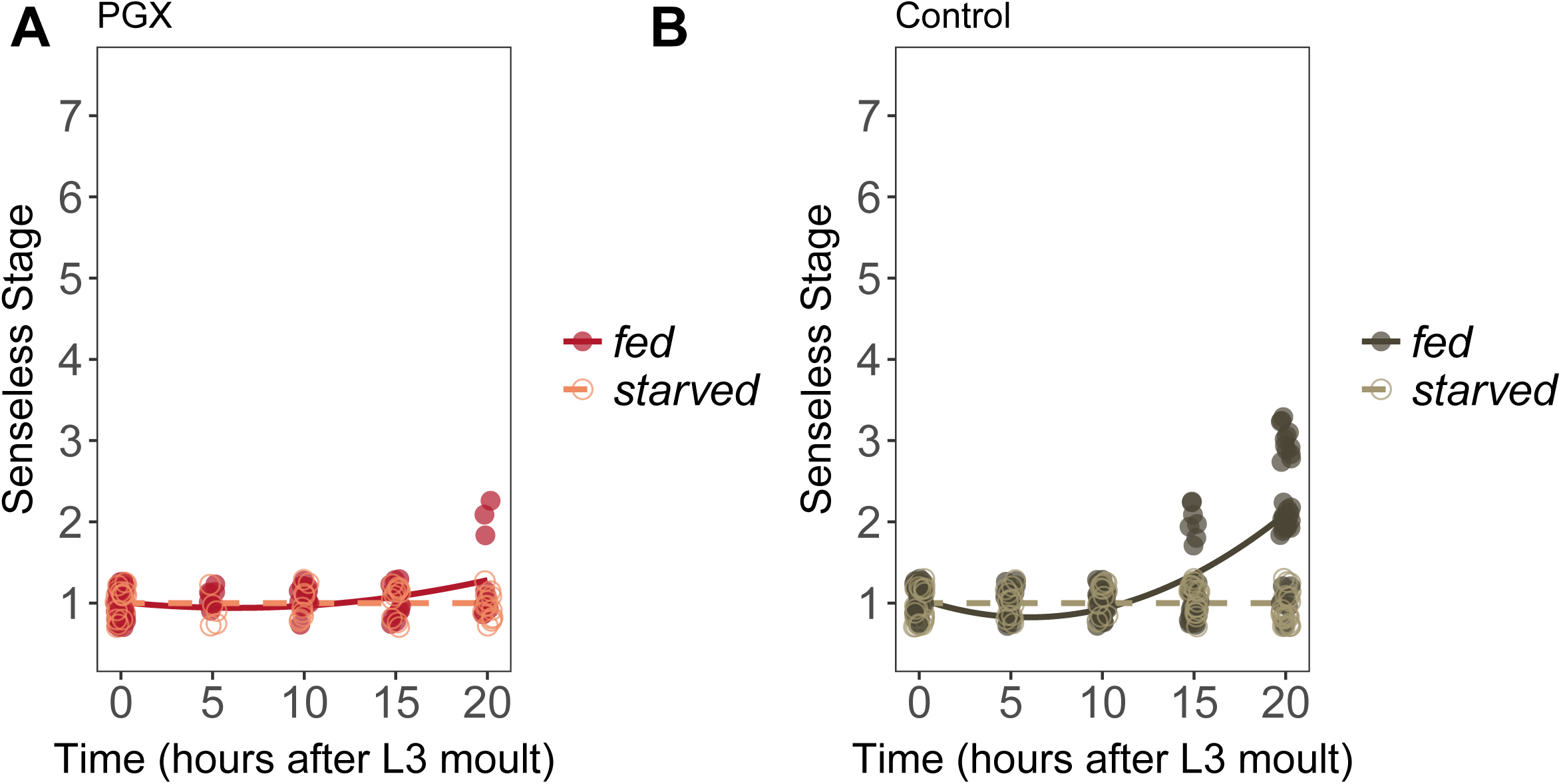
Senseless patterning in wing discs from fed and starved PGX and control larvae. (A) Pa’erning does not progress in either fed or starved PGX larvae. (B) Pa’erning does not progress in starved control larvae but does in fed control larvae. There is a significant interaction between the effects of time and food on Achaete pa’erning in control larvae (linear regression: *F*_food*time_ =67.98, *P* < 0.001). In PGX larvae Senseless pa’erning does not progress at all in starved larvae (linear regression: *F*_time_ =0.057, *P* =0.82), but does in fed larvae (linear regression: *F*_time_ =9.76, *P* < 0.01). Control genotypes are the pooled results from both parental controls (either the *phm-GAL4; GAL80ts* or *UAS-GRIM* parental strain crossed to w^1118^). Each point corresponds to a wing disc, N_PGX - starved_ = 67, N_PGX - fed_ = 74, N_control - starved_ = 118, N_control - fed_ = 151 across all time points.

