## Supplemental Figures and Tables for "Ecdysone coordinates plastic growth with robust pattern in the developing wing"

#### Supplementary Figures

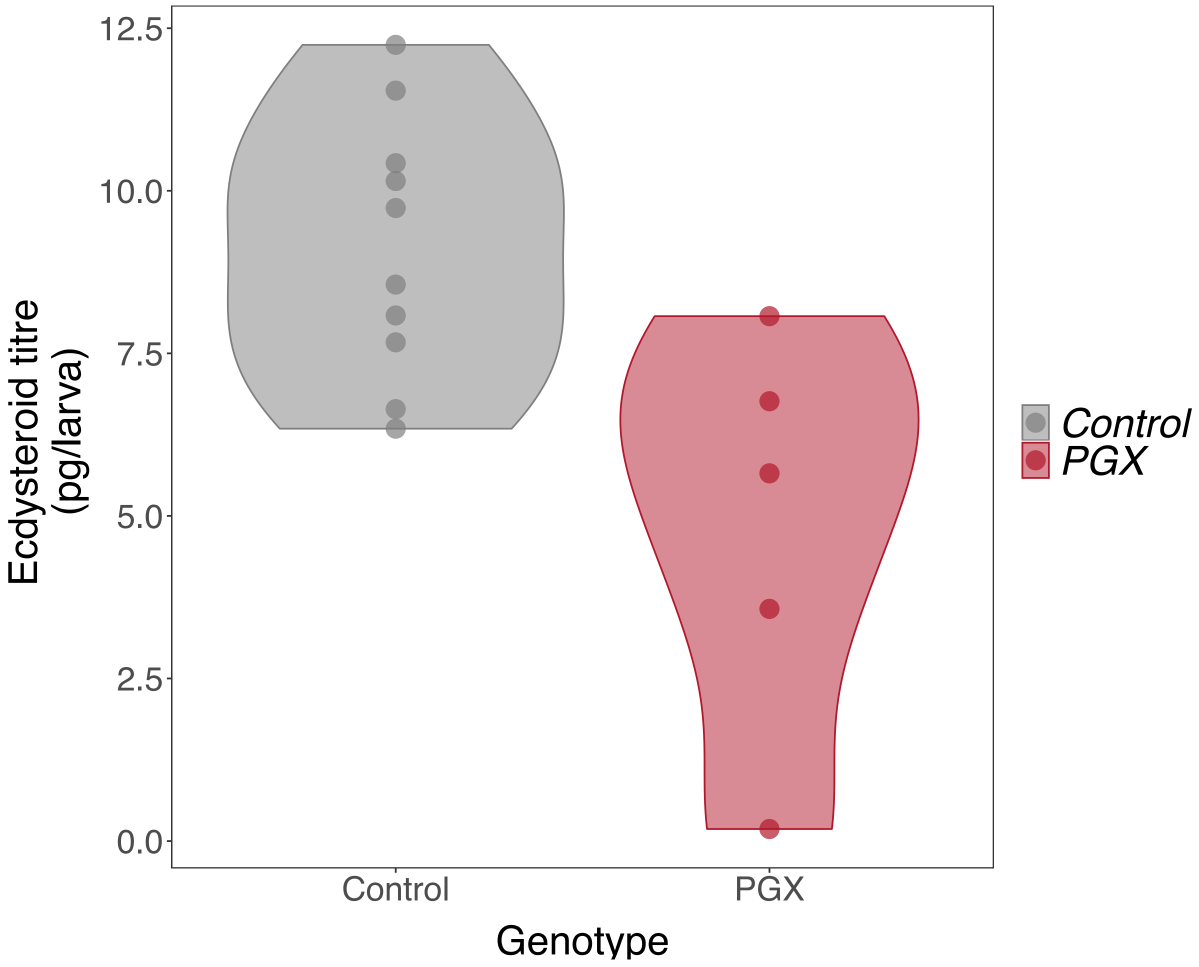


**Figure 2 Supplement 1:** Ecdysteroid titres in PGX and control larvae. Newly ecdysed larvae were placed on sucrose/yeast diets. Ecdysteroid titres in control (phm > + and + > grim) are significantly higher ecdysteroid titres than PGX larvae ,as determined by linear models and pairwise comparisons of the means (F $value_{1, 13}$ = 10.75, p value = 0.006, $N_{control}$ = 10, $N_{PGX}$ = 5). Data is plotted with violin plots and the individual replicates (5 per genotype) are included as points overlaying the violin plots.

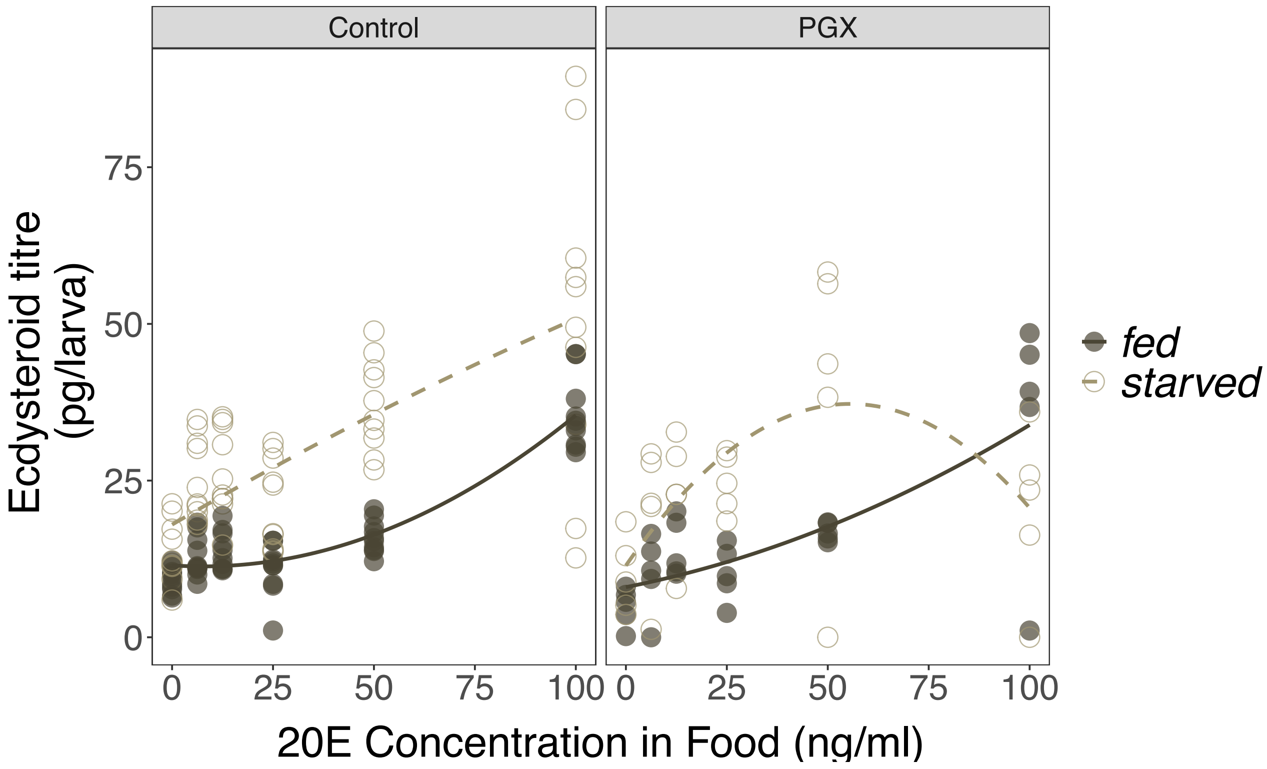


**Figure 7 Supplement 1:** The effects of 20E concentration in the food on ecdysteroid titres in control and PGX larvae. Newly ecdysed larvae were placed on sucrose/yeast (fed)or 20% sucrose/1% agar (starved) diets supplemented with a range of 20E concentrations for 20h. They were then transferred to the same diet without 20E and dyed with blue food colouring for 2 h, to eliminate residual 20E in the gut. Ecdysteroid titres in fed and starved control (phm > + and + > grim) and PGX larvae fed a range of 20E concentrations. There is a significant positive relationship between 20E concentration in the food and the concentration of ecdysteroids in the larvae, as indicated by a significant 20E term (Supplementary Table 8). Furthermore, starved larvae had higher ecdysone titres than fed larvae. Open and closed points represent the biological replicates, and solid and dashed lines are linear regressions. $N_{control - starved}$ = 60, $N_{control - fed}$ = 60, $N_{PGX - starved}$ = 30, $N_{PGX - fed}$ = 30 across all 20E concentrations.

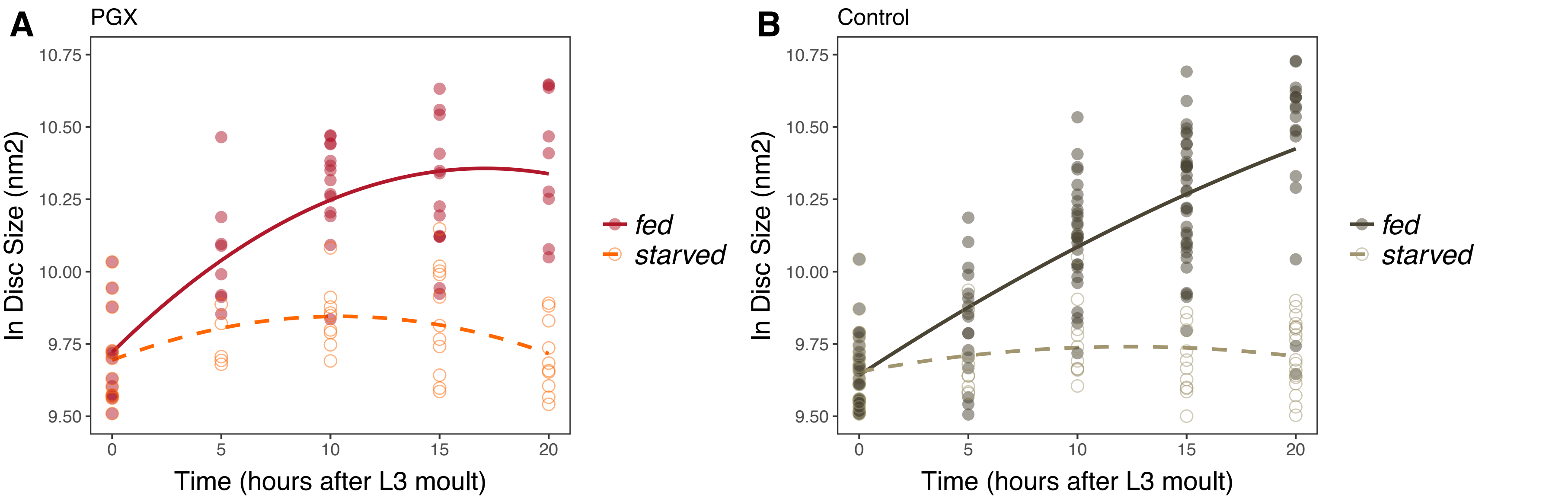


**Figure 7 Supplement 2:** Wing imaginal disc growth is suppressed in fed PGX larvae relative to controls, and in starved larvae of both genotypes. Wing disc growth was modelled as a quadratic, and there was a significant interaction between genotype (PGX v. Control) and nutrition (fed v. starved) on growth (Supplementary Table 7). Solid line/closed point = fed larvae, broken line/open point = starved larvae. Each point corresponds to a wing disc, $N_{PGX\text{ - starved}}$ = 67, $N_{PGX\text{ - fed}}$ = 74, $N_{control\text{ - starved}}$ = 118, $N_{control\text{ - fed}}$ = 151 across all time points.

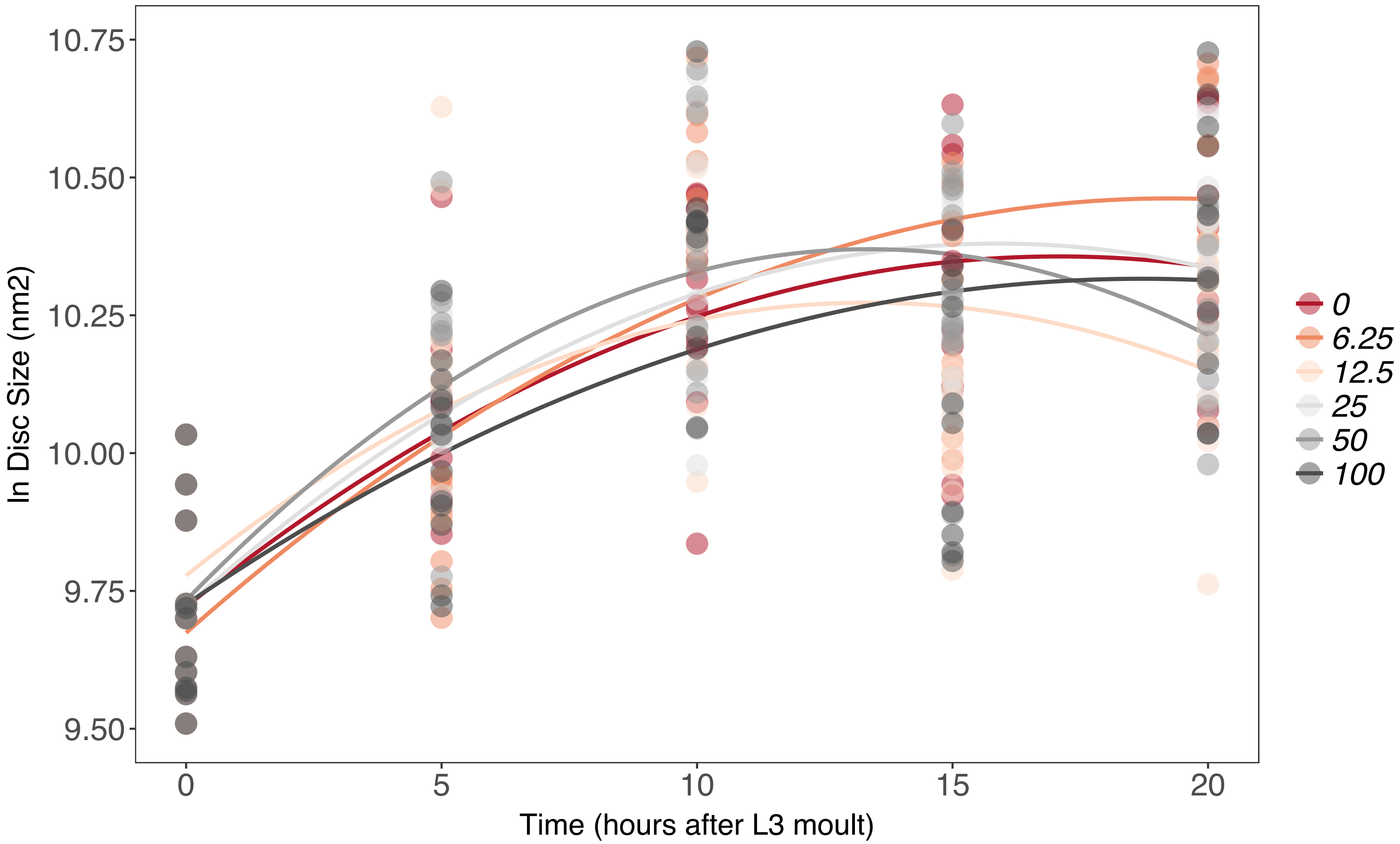


**Figure 7 Supplement 3:** There is no effect of supplemental 20E on growth of the wing imaginal disc in fed PGX larvae. Growth was modelled as $S = E + T + T^2 + E*T + E*T^2$, where $S$ = disc size, $E$ = 20E concentration, and $T$ = disc age. There was no significant effect of $E$ on the linear or quadratic growth rate of the wing imaginal discs (Supplementary Table 8). Each point corresponds to a wing disc, $N_{PGX-fed}$ = 459 (73-86 discs were sampled per treatment across all time points).

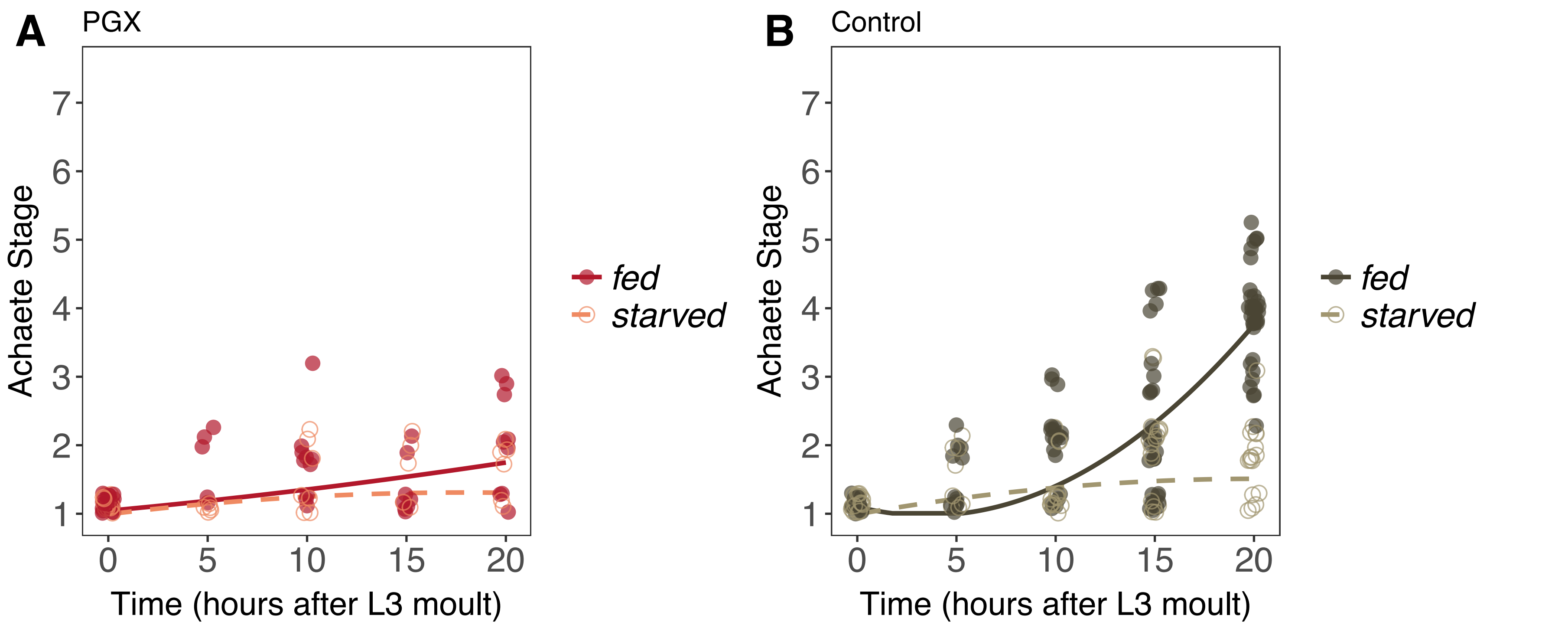


**Figure 8 Supplement 1** Achaete patterning in wing discs from fed and starved PGX and control larvae. (A) Patterning does not progress in either fed or starved PGX larvae. (B) Patterning does not progress in starved control larvae but does in fed control larvae. There is a significant interaction between the effects of disc age and food on Achaete patterning in control larvae (orthogonal polynomial regression: $F_{food*disc\ age^2}$=67.98, P < 0.001), but not in PGX larvae (orthogonal polynomial regression: $F_{food*disc\ age^2}$=1.81, P = 0.163). Each point corresponds to a wing disc, $N_{PGX\text{ - }starved}$ = 67, $N_{PGX\text{ - }fed}$ = 74, $N_{control\text{ - }starved}$ = 118, $N_{control\text{ - }fed}$ = 151 across all time points.

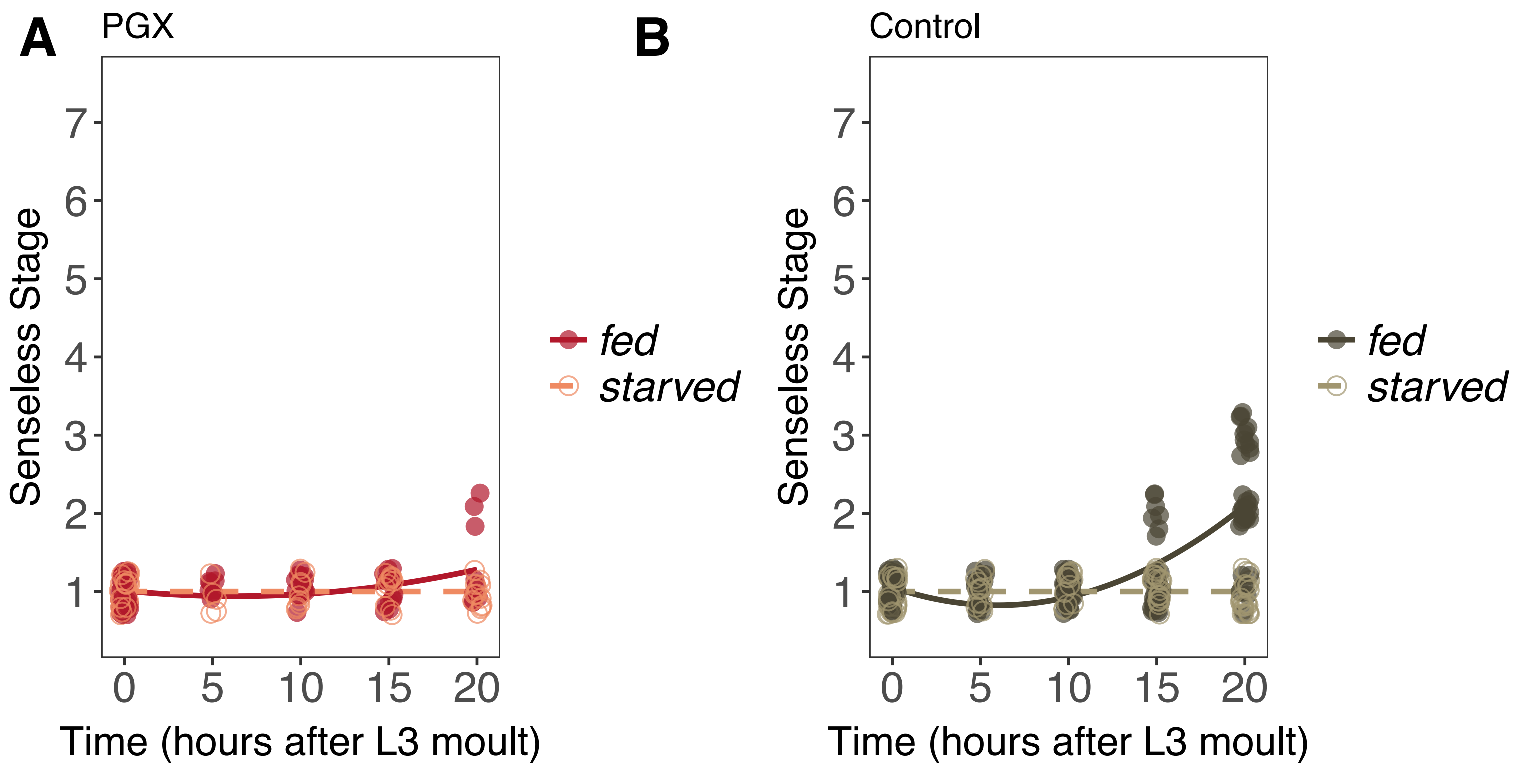


**Figure 9 Supplement 1**: Senseless patterning in wing discs from fed and starved PGX and control larvae. (A) Patterning does not progress in either fed or starved PGX larvae. (B) Patterning does not progress in starved control larvae but does in fed control larvae. There is a significant interaction between the effects of time and food on Achaete patterning in control larvae (linear regression: $F_{\mathrm{food*time}}$=67.98, $P < 0.001$). In PGX larvae Senseless patterning does not progress at all in starved larvae (linear regression: $F_{\mathrm{time}}$=0.057, $P$ =0.82), but does in fed larvae (linear regression: $F_{\mathrm{time}}$=9.76, $P < 0.01$). Control genotypes are the pooled results from both parental controls (either the *phm-GAL4; GAL80ts* or *UAS-GRIM* parental strain crossed to $w^{1118}$). Each point corresponds to a wing disc, $N_{\mathrm{PGX\text{ - }starved}}$ = 67, $N_{\mathrm{PGX\text{ - }fed}}$ = 74, $N_{\mathrm{control\text{ - }starved}}$ = 118, $N_{\mathrm{control\text{ - }fed}}$ = 151 across all time points.

### Supplementary Tables

**Supplementary Table 1:** Comparing the Gompertz growth curve parameters for wing imaginal discs from PGX and control (+>Grim and phm>+) larvae.

| Genotype/Condition | Asymptote (*a*)[A] | *F*[B] | Displacement along x (*b*)[A] | *F*[B] | Growth rate (*g*)[A] | *F*[B] |
|---|---|---|---|---|---|---|
| PGX | 10.53 (10.45–10.65) | 52.62*** | 0.11 (0.10–0.13) | 68.91*** | 0.90 (0.85–0.94) | 5.45* |
| Control | 11.57 (11.42-11.74) | | 0.22 (0.21–0.24) | | 0.95 (0.94–0.95) | |

[A] Parameter values are for a Gompertz model $y = a \cdot e^{-b \cdot e^{-g \cdot x}}$,where x is disc size and y is Achaete pattern. Values in parentheses are 95% confidence intervals.
[B] *F*-test of whether parameter value differs between genotype/condition. * *P* value < 0.05, *** *P* value < 0.0001.

**Supplementary Table 2:** Comparing the Gompertz curve parameters of Achaete patterning against time for wing imaginal discs from PGX and control (+>Grim and phm>+) larvae.

| Genotype/Condition | Asymptote (*a*)[A] | *F*[B] | Displacement along x (*b*)[A] | *F*[B] | Growth rate (*g*)[A] | *F*[B] |
|---|---|---|---|---|---|---|
| PGX | 3.98 (3.56–4.41) | 12.62*** | 1.03(0.84–1.21) | 12.05*** | 0.045 (0.029-0.063) | 0.56 |
| Control | 6.34 (6.20–7.08) | | 1.64 (1.42–1.87) | | 0.074 (0.059-0.090) | |

[A] Parameter values are for a Gompertz model $y = a \cdot e^{-b \cdot e^{-g \cdot x}}$,where x is disc size and y is Achaete pattern. Values in parentheses are 95% confidence intervals.
[B] *F*-test of whether parameter value differs between genotype/condition. * *P* value < 0.05, *** *P* value < 0.0001.

**Supplementary Table 3:** Comparing the linear Senseless patterning parameters for wing imaginal discs from PGX and control (+>Grim and phm>+) larvae.

| Genotype | Intercept (*a*)[A] | F[B] | Slope (*b*)[A] | F[B] |
|---|---|---|---|---|
| PGX | 1.25 (0.94–1.55) | 25.89*** | 0.01(0.00–0.02) | 583.26*** |
| Control | 0.46 (0.21–0.72) | | 0.13 (10.12–0.14) | |

[A] Parameter values are for a linear model $y = B + Ax$, where x is disc size and y is Senseless pattern. Values in parentheses are 95% confidence intervals.
[B] *F*-test of whether parameter value differs between genotype/condition. * *P* value < 0.05, *** *P* value < 0.0001.

**Supplementary Table 4:** The effect of supplementing 20-hydroxyecdysone (20E) on wing disc size, Achaete stage, and Senseless stage in control and PGX larvae

| Factors | F value – Disc Size | F value – Achaete Stage | F value – Senseless Stage |
|---|---|---|---|
| Genotype[A] | 162.78*** | 210.49*** | 265.06*** |
| 20E Treatment[B] | 135.95*** | 56.38*** | 129.21*** |
| Genotype * Treatment | 72.87*** | 120.26*** | 163.72*** |

[A] Genotypes include PGX and control larvae.
[B] 20E Treatment includes supplementation with 0.15 mg/ml of 20E or the same volume of ethanol
*** *P* value < 0.001

**Supplementary Table 5:** The effect of supplementing 20-hydroxyecdysone (20E) on wing disc size, Achaete stage, and Senseless stage in control and PGX larvae reared on starvation medium or on fly food.

| Factors | F value – Disc Size | F value – Achaete Stage | F value – Senseless Stage |
|---|---|---|---|
| Genotype[A] | 1.15 | 0.48 | 0.00 |
| 20E + Diet Treatment[B] | 214.20*** | 358.80*** | 117.22*** |
| Genotype * 20E + Diet Treatment | 28.86*** | 99.76*** | 72.08*** |

[A] Genotypes include PGX and control larvae.

[B] 20E + Diet Treatment includes starved + 0.15 mg/ml 20E, starved + ethanol, and fed

*** $P$ value < 0.001

**Supplementary Table 6:** Comparing the parameters of the linear relationship for Achaete pattern against (log) disc size, between phm > InR and their parental control line (+ > InR), and between P0206 > PTEN and their parental control line (+ > PTEN).

| Genotype/ Condition | Intercept ($a$) [A] | $F$[B] | Slope ($b$) [A] | $F$[B] |
|---|---|---|---|---|
| phm>InR | -14.99 (-17.32–-12.65) | | 1.93 (1.70-2.15) | |
| P0206>PTEN | - 12.23(-13.72–-10.75) | 22.42*** | 1.61 (1.44–1.78) | 660.67** |
| Control | -16.40 (-18.45–-14.23) | | 1.97 (1.83-2.10) | |

[A] Parameter values are for a linear model $y = B + Ax$, where x is disc size and y is Achaete pattern. Values in parentheses are 95% confidence intervals.

[B] $F$-test for when the parameter value differs between genotype/condition. [NS] $P$ value > 0.05, *$P$ value < 0.05, *$P$ value < 0.01, *** $P$ value < 0.001.

**Supplementary Table 7:** Comparing the parameters of the logistic relationship for Senseless pattern against (log) disc size, between Samarkand larvae reared at 18°C, 25°C, and 29°C, between phm > InR and their parental control line (+ > InR), and between P0206 > PTEN and their parental control line (+ > PTEN).

| Genotype/ Condition | Minimum (*a*)[A] | F[B] | Maximum (*b*)[A] | F[B] | Point of inflection (*c*)[A] | F[B] | Logistic Growth rate (*d*)[A] | F[B] |
|---|---|---|---|---|---|---|---|---|
| Phm>InR | 0.92 (0.32 – 1.52) | 0.12[NS] | 7.13 (5.72–8.54) | 3.61* | 10.42 (10.14–10.69) | 6.94** | 2.13 (1.40 – 4.45) | 5.89** |
| P0206>PTEN | 1.00 (0.73-1.27) | | 7.15 (6.31–8.00) | | 10.97 (10.84–11..10) | | 4.65 (3.31–7.84) | |
| Control | 1.04 (0.80-1.29) | | 8.91 (7.08–10.73) | | 11.22 (10.99–11.45) | | 2.39 (1.86–3.35) | |

[A] Parameter values for the four-parameter logistic model $y = a + (b - a)/\left(1 + e^{d(c-x)}\right)$ where x is disc size and y is Senseless pattern. Values in parentheses are Bonferroni-corrected 95% confidence intervals.

[B] *F*-test for when the parameter value differs between genotype/condition. [NS] *P* value > 0.05, **P* value < 0.05, **P* value < 0.01, *** *P* value < 0.001.

**Supplementary Table 8:** Effect of 20E supplementation on titres of ecdysteroids in control (phm > + and + > grim) and PGX larvae under starved and fed conditions.

| Factor | df | F value | *P* |
|---|---|---|---|
| 20E (poly, 2) | 1 | 69.60 | <0.001 |
| Food | 2 | 59.14 | <0.001 |
| Genotype Group | 1 | 6.05 | 0.015 |
| 20E (poly, 2): Food | 1 | 6.14 | 0.003 |
| 20E (poly, 2): Genotype Group | 2 | 7.32 | 0.001 |
| Food: Genotype Group | 2 | 2.53 | 0.113 |
| 20E (poly, 2): Food: Genotype Group | 2 | 6.21 | 0.003 |

**Supplementary Table 9:** Effect of diet type on wing disc growth in PGX and control (phm > + and + > grim) larvae under starved and fed conditions.

| Factor | df | *F* value | *P* |
|---|---|---|---|
| Genotype | 1 | 1.4 | 0.24 |
| Food | 1 | 278.5 | <0.001 |
| $Time^2$ | 2 | 202.5 | <0.001 |
| Genotype: Food | 1 | 8.7 | 0.003 |
| Genotype: $Time^2$ | 2 | 13.2 | <0.001 |
| Food: $Time^2$ | 2 | 98.7 | <0.001 |
| Genotype: Food: $Time^2$ | 2 | 5.31 | 0.005 |

**Supplementary Table 10:** Effect of 20E concentration on growth of the wing imaginal disc in starved and fed PGX larvae.

| | Starved | | | | Fed | | | |
|---|---|---|---|---|---|---|---|---|
| Factor[A] | SS | df | *F* | *P* | SS | df | *F* | *P* |
| 20E | 1.5 | 1 | 27.3 | <0.001 | 0 | 1 | 1.35E-01 | 0.7137 |
| Disc Age$^2$ | 7.2 | 2 | 64.6 | <0.001 | 36.6 | 2 | 29.8 | <0.001 |
| 20E: Disc Age$^2$ | 1 | 2 | 8.88 | <0.001 | 0 | 2 | 4.01E-01 | 0.6697 |
| Residuals | 22.3 | 403 | | | 27.8 | 453 | | |

[A] Factors were fit using an orthogonal polynomial regression.

**Supplementary Table 11:** Comparisons of model fit for linear rates of wing disc growth, Achaete patterning, and Senseless patterning across 20E concentrations in fed and starved larvae.

.

| | | Wing growth rate - starved | | Achaete patterning rate - fed | | Achaete patterning rate - starved | | Senseless patterning rate - fed | | Senseless patterning rate - fed | |
|---|---|---|---|---|---|---|---|---|---|---|---|
| | df | AIC | BIC | AIC | BIC | AIC | BIC | AIC | BIC | AIC | BIC |
| Michaelis Menten[A] | 4 | **-43.67** | **-44.50** | -22.72 | -23.55 | -18.69 | -19.52 | -29.88 | -30.71 | -29.64 | -30.48 |
| Three parameter logistic[B] | 4 | -30.72 | -31.55 | -25.28 | -26.12 | -29.83 | -30.66 | -40.84 | -41.68 | **-35.09** | **-35.93** |
| Four parameter logistic[C] | 5 | -41.73 | -42.77 | **-32.99** | **-34.03** | **-46.67** | **-47.71** | **-48.52** | **-49.56** | -33.37 | -34.41 |

[A] Michaelis Menten function: $y = c + (d-c)/(1+(b/x))$, where *c* is *y* at *x*=0, *d* = *y[max]*, and *b* is *x* where *y* is halfway between *c* and *d*.

[B] Three-parameter log-logistic function: $y = d/(1+e^{(b(\log(x)-\log(a))})$, where *d* = *y[max]*, *b* is the rate of increase, and *a* is the inflection point. Note, in the three-parameter log-logistic function, c = 0.

[C] Four-parameter log-logistic function: $y = c + (d-c)/(1+e^{(b(\log(x)-\log(a))})$, where *c* is *y* at *x*=0, *d* = *y[max]*, *b* is the rate of increase, and *a* is the inflection point.

The selected model has the lowest AIC and BIC values, and is in bold.

**Supplementary Table 12:** Effect of 20E concentration on Achaete patterning stage in the wing imaginal disc in fed and starved PGX larvae.

| | Fed | | | | Starved | | | |
|---|---|---|---|---|---|---|---|---|
| Factor[A] | SS | df | *F* | *P* | SS | df | *F* | *P* |
| 20E | 124.9 | 5 | 72.65 | <0.001 | 152.7 | 5 | 75.82 | <0.001 |
| Disc Age$^2$ | 4.7 | 2 | 6.78 | 0.0013 | 1.4 | 2 | 1.70 | 0.18 |
| 20E: Disc Age$^2$ | 108.3 | 10 | 31.48 | <0.001 | 178.0 | 10 | 44.20 | <0.001 |
| Residuals | 151.7 | 441 | | | 157.48 | 391 | | |

[A] Factors were fit using an orthogonal polynomial regression.

**Supplementary Table 13:** Effect of 20E concentration on Senseless patterning stage in the wing imaginal disc in fed and starved PGX larvae.

| | Fed | | | | Starved | | | |
|---|---|---|---|---|---|---|---|---|
| Factor[A] | SS | df | *F* | *P* | SS | df | *F* | *P* |
| 20E | 34.3 | 5 | 44.55 | <0.001 | 56.7 | 5 | 55.08 | <0.001 |
| Disc Age$^2$ | 0.72 | 2 | 2.34 | 0.097 | 0 | 2 | 0 | 1 |
| 20E: Disc Age$^2$ | 37.4 | 10 | 24.28 | <0.001 | 70.45 | 10 | 27.36 | <0.001 |
| Residuals | 67.84 | 441 | | | 100.7 | 391 | | |

[A] Factors were fit using an orthogonal polynomial regression.
